# Ligands tune the local and global motions of neurotensin receptor 1 (NTS1): a DFT-guided solution NMR analysis

**DOI:** 10.1101/2022.08.09.503369

**Authors:** Fabian Bumbak, Miquel Pons, Asuka Inoue, Juan Carlos Paniagua, Fei Yan, Hongwei Wu, Scott A. Robson, Ross A. D. Bathgate, Daniel J. Scott, Paul R. Gooley, Joshua J. Ziarek

## Abstract

Unlike many signaling proteins that function as binary switches between ‘on and off’ states, G protein-coupled receptors (GPCRs) exhibit basal activity that can be increased or decreased by numerous ligands. A given receptor can recognize multiple ligands, allosteric modulators, and transducers to create a complex free energy landscape. Many of the lowest energy states have been captured by static structural techniques while detailing the wells’ widths, metastable states, and the transition between them, is still in its infancy. Nuclear magnetic resonance (NMR) spectroscopy can monitor the structure and dynamics of GPCR ensembles across fifteen orders-of-magnitude, but technical challenges have limited its application to super-microsecond timescales. Focusing on a prototypical peptide-binding GPCR, the neurotensin receptor 1 (NTS_1_), we employed NMR and density functional theory (DFT) to probe global sub-nanosecond motions. The near random coil chemical shifts of the apo receptor produced a poor correlation with theoretical predictions that may indicate a high degree of conformational averaging in solution, a crystallization artifact, or both. Whereas orthosteric agonists and antagonists both rigidified the receptor, but to varying degrees, which suggests conformational entropy differentially contributes to their respective pharmacology. The strong correlations of observed and theoretical chemical shifts lend confidence to interpreting spectra in terms of local structure, methyl dihedral angle geometry, and pico-second timescale transitions. Together, our results suggest a role for sub-nanosecond dynamics and conformational entropy in GPCR ligand discrimination.

## INTRODUCTION

G protein-coupled receptors (GPCRs) are the largest family of membrane proteins – comprising approximately 3% of the human genome. They recognize a diverse set of stimuli at the plasma membrane to regulate processes from vision, smell and taste, to immune, neurologic and reproductive functions. As fundamental components of all major systems it is no surprise that GPCRs dominate the therapeutic market – accounting for more than 30% of FDA-approved drugs ^1-2^. Activation is initiated by agonist binding on the extracellular face, which produces a conformational change on the intracellular side. Unlike many signaling proteins that function as binary switches between ‘on and off’ states, GPCRs feature a ligand-independent basal activity that is increased or decreased upon ligand binding, and then further regulated by allosteric modulators. Activated receptors signal intracellularly through G protein and arrestin transducers equally (balanced signaling) or selectively (biased signaling). A single receptor may specifically recognize several ligands and respond uniquely to each, creating a complex conformational landscape. Thus, there is immense therapeutic potential in the ability to tune receptor signaling using partial agonists, biased agonists, and allosteric modulators, that is only beginning to be tapped ^3-4^.

The principles of allostery, whereby ligand association at one site elicits altered activity at a remote location, have been critical to our understanding of GPCR signaling ^5-8^. A dramatic example from crystallography is that orthosteric ligand binding translates the intracellular portions of transmembrane (TM) helices 5 and 6, which are located > 40 Å away, outward to accommodate heterotrimeric G proteins ^9^. The Monod-Wyman-Changeux (MWC) model of allostery (i.e. conformational selection) posits the pre-existence of both inactive and active conformations whose relative populations are modulated by the ligand ^10^. Indeed, elegant nuclear magnetic resonance (NMR) spectroscopy studies using ^19^F-labeled TMs confirm a conformational ensemble that in most instances includes more than two stable, low-energy states ^11^. Yet, in many instances the MWC model is unsatisfactory for understanding the full pharmacological landscape of partial agonists, allosteric modulators, and biased agonists ^7-8, 11^. There is a growing body of evidence that allostery doesn’t necessarily require conformational changes on a scale that can be observed by static structural techniques such as cryo-EM and X-ray crystallography. First theorized by Cooper and Dryden nearly 35 years ago^12^, dynamically-driven (DD) allostery asserts that the frequency and amplitude of sub-nanosecond motions around the average conformation (i.e. conformational entropy) can effectively reduce the energy barrier between inactive-active modes without the need for structural change.

NMR spectroscopy is uniquely capable of reporting on both types of allostery at near atomic resolution but, to date, has primarily focused on the role of MWC allostery in GPCR systems. This is largely due to the technical challenges of selective isotope labeling in eukaryotic cells and the resonance assignment-by-mutagenesis approach^11^. Although there are examples that demonstrate a role for sub-nanosecond backbone dynamics in protein function^13^, the fast motions of methyl-bearing side chain are clearly more informative reporters on DD allostery. Quantification of local sidechain motions, in the form of NMR generalized order parameters, have been used to develop an empirical, model-independent proxy for conformational entropy ^14-16^. The comparatively high mobility of methionine side chains, and low sequence abundance, has generally diminished their inclusion in these motional analyses. Yet, for practical reasons, selective methionine labeling remains one of the most commonly applied approaches to studying GPCRs by NMR ^11^.

Determination of the generalized order parameter is a far more laborious process than that of other NMR observables such as the chemical shift and linewidths. Chemical shift values reflect a nucleus’ local electronic environment, which makes them extremely sensitive probes of both structure and dynamics; for example, the ^13^C methyl chemical shift of branched-chain amino acids (e.g. Ile, Leu, Val) is strongly influenced by the side-chain rotameric state (trans, gauche) ^17-19^. Chashmniam and colleagues recently used density functional theory (DFT) quantum calculations to demonstrate that the (de-)shielding effect of neighboring atoms on the methionine methyl chemical shift is comparable, or greater, than the one arising from local side-chain geometry ^20^. They showed that the relative ^13^C chemical shift values of methionine methyl groups can be predicted from a static high-resolution crystal structure by only considering atoms within a 6 Å sphere. When a linear correlation is observed between the theoretical and experimental ^13^C chemical shift values of methionines located far apart in the structure, it indicates a similar degree of side-chain conformational averaging throughout the protein. The (common) degree of local conformational averaging is protein dependent and can be extracted from the slope of the regression line. Thus, this slope can be interpreted as an order parameter for the global flexibility of the protein, as detected locally by multiple methionine side chains in distant parts of the structure. This methionine chemical shift global order parameter (S_MCS_) can theoretically range from one (completely rigid) to zero (completely averaged). In other words, the experimental chemical shifts are scaled from the rigid-structure calculated values toward those expected from a totally flexible environment.

Here, we focus on a prototypical peptide-binding receptor, neurotensin receptor 1 (NTS_1_) to test if S_MCS_ can sense ligand-dependent changes in global receptor dynamics. We functionally validated and assigned the ^13^C^ε^H_3_-methionine resonances of a minimal-methionine NTS_1_ variant in the presence of several orthosteric and allosteric ligands. We then compared the experimental ^13^C^ε^ chemical shifts to DFT-calculated chemical shifts from high resolution apo, agonist and antagonist bound NTS_1_ crystal structures ^21^. Whereas the basal ensemble of the apo receptor appears highly dynamic, agonist and antagonist binding tune global motions that differentially modulate receptor rigidity. The strong linear correlation between experimental chemical shifts and DFT calculations suggested that mechanistic hypotheses could be derived from detailed structural examination. Given the number of ^13^C^ε^H_3_-methionine NMR studies ^11^ and corresponding high-resolution GPCR crystal structures, our results reveal a tractable approach to exploring global receptor dynamics on the sub-nanosecond timescale.

## RESULTS

### NTS_1_ construct design and ^13^C^ε^H_3_-methionine chemical shift assignment

Our study focuses on a selectively ^13^C^ε^H_3_-methionine labelled NTS_1_ construct, termed enNTS_1_ΔM4, which was derived from a previously thermostabilized rNTS_1_ variant (enNTS_1_) ^22^ by removing four (M181L, M267L, M293L and M408V) of the ten endogenous methionine residues (Figure 1A). Preliminary experiments showed that mutagenesis did not adversely affect structural integrity as it had little to no observable effect on the remaining resonances’ chemical shifts (Figure S1). M267^5.68^, M293^ICL3^ and M408^H8^ (superscript refers to Ballesteros-Weinstein numbering ^23^) are solvent exposed and highly degenerate in two-dimensional (2D) ^1^H-^13^C heteronuclear multiple quantum correlation (HMQC) spectra while M181^ICL2^ overlaps with M352^7.36^ in some instances. Thus, the mutations simplify the analysis of 2D ^1^H-^13^C HMQC spectra (Figure S1), while preserving the structure of enNTS_1_. enNTS_1_ΔM4 retains six endogenous methionine residues (M204^4.60^, M208^4.64^, M244^5.45^, M250^5.51^, M330^6.57^ and M352^7.36^) (Figure 1A) with all, except for M244^5.45^ being retained across species. Four of these methionines (M204^4.60^, M208^4.64^, M330^6.57^ and M352^7.36^) are also conserved in all NTS_2_ sequences, but M250^5.51^ is the only probe significantly conserved among peptide GPCRs (19%, ranking 2nd after leucine).

**Figure 1.**
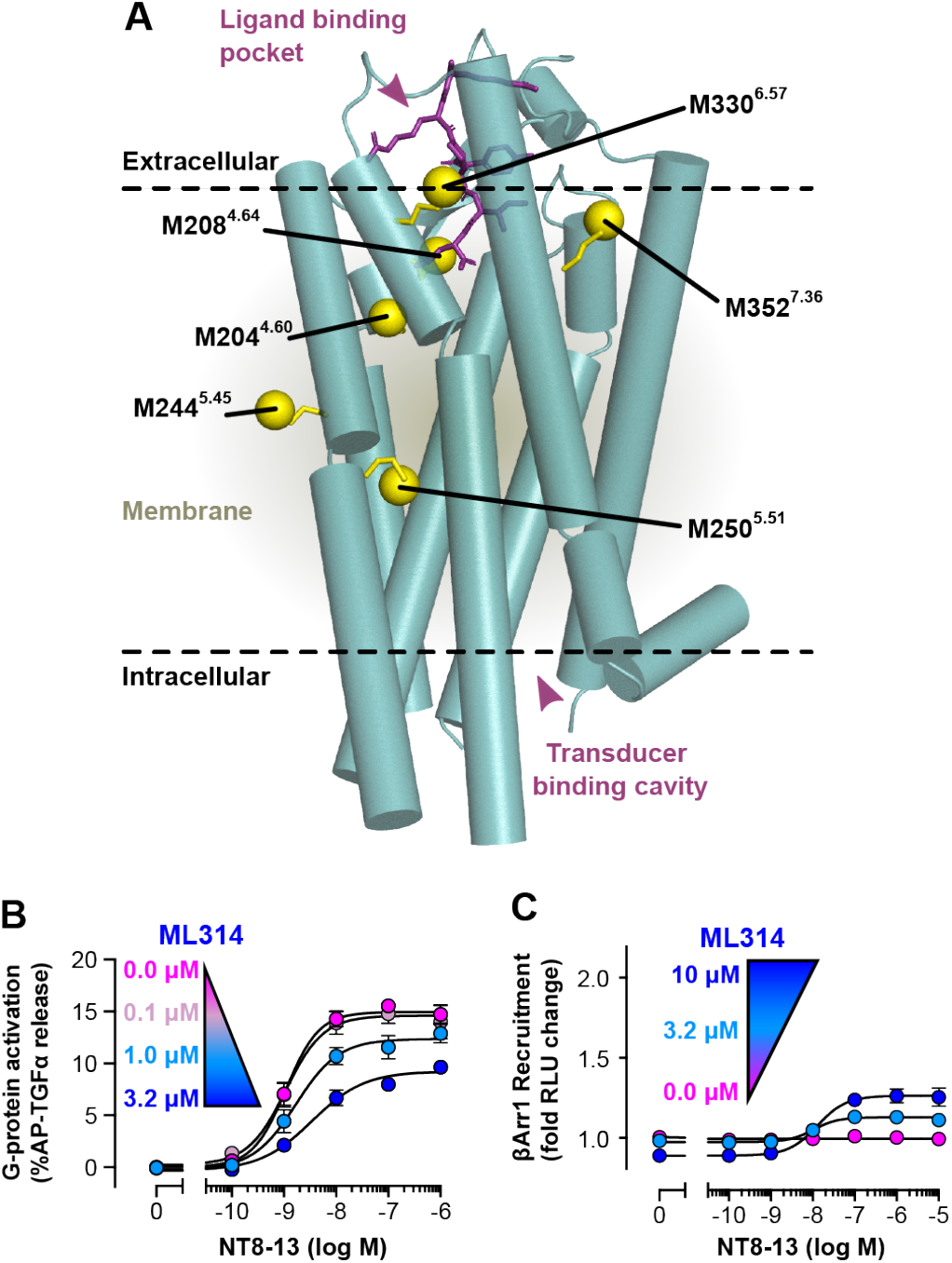
Ligand-induced chemical shift changes observed for ^13^CH_3_-methionine labelled enNTS_1_ΔM4. A) Cylindrical representation of thermostabilized rNTS_1_ (PDB 4BWB) with labelled methionine methyl groups shown as yellow spheres (superscript - Ballesteros-Weinstein nomenclature ^23^) and NT8-13 shown as purple sticks. B) ML314 reduces efficacy of NT8-13 dependent, enNTS_1_ΔM4-mediated G protein activation in a TGFα shedding assay using HEK293A cells ^24^. Cells were stimulated with increasing concentrations of NT8-13 in the absence (magenta) or presence of 0.1 μM (light pink), 3.2 μM (light blue), and 10 μM (blue) ML314. Error bars represent SEM from three independent experiments. C) ML314 potentiates NT8-13 dependent βArr1 recruitment to enNTS_1_ΔM4. βArr1 recruitment was measured by a NanoBiT-based assay using HEK293A cells ^75^. Cells were stimulated with increasing concentrations of NT8-13 in the absence (magenta) or presence of 3.2 μM (light blue), and 10 μM (blue) ML314. Note that there was a modest β-arrestin suppressive effect with 10 μM ML314 alone (dark blue symbol at 0 μM NT8-13) yet showing response to NT8-13 in a concentration-dependent manner. Error bars represent SEM from three independent experiments.

The functional integrity of enNTS_1_ΔM4 was assessed using a cell-based alkaline phosphatase (AP) reporter assay for G protein activation. Stimulation of Gα_q_ and Gα_12/13_ leads to ectodomain shedding of an AP-fused transforming growth factor-α (TGFα), which is then quantified using a colorimetric reporter ^24^. HEK293A cells were transfected with AP-TGFα and a NTS_1_ plasmid construct. NT8-13, the smallest fully functional fragment corresponding to residues 8-13 of NT ^25^, stimulates robust, concentration-dependent G protein-coupling to enNTS_1_ΔM4 in the TGFα shedding assay (Figures 1B and S2A). ML314 attenuates agonist-mediated G protein activation of enNTS_1_ΔM4 in a concentration-dependent manner (Figures 1B and S2B) as previously reported ^26-27^. βArr1 recruitment was measured using a NanoBiT enzyme complementation system ^28^. The large and small fragments of the split luciferase were fused to the N-terminus of βArr1 and the C-termini of NTS_1_ variants, respectively, and these constructs were expressed in HEK293A cells. As a negative control, we used the vasopressin V2 receptor (V2R) C-terminally fused with the small luciferase fragment. The basal βArr1 recruitment of enNTS_1_ΔM4 did not increase upon agonist addition (Figures 1C and S2C) but, similar to previous observations with βArr2, ML314 potentiates NT8-13-mediated βArr1 recruitment (Figure 1C). Although ML314 was previously reported to stimulate βArr1 recruitment independent of NT8-13 ^27^, we only observe βArr-biased allosteric modulator (BAM) pharmacology (Figure S2D). It is possible that the agonist pharmacology of ML314 may require even higher ligand concentrations which are incompatible with the NanoBiT assay. Both enNTS_1_ΔM4 and hNTS_1_ were equally expressed on the cell surface (Figure S2E).

enNTS_1_ΔM4 resonances were assigned using a “knock-in” strategy starting with the binding-competent, minimal methionine enNTS_1_ΔM8 mutant containing only M330^6.57^ and M352^7.36^. For this approach, six plasmids were engineered with sequentially reintroduced methionine residues. All enNTS_1_ variants were expressed in *Escherichia coli* by inhibiting the methionine biosynthesis pathway while supplementing a defined medium with ^13^C^ε^H_3_-methionine. Final NMR samples were prepared in n-dodecyl-β-D-maltopyranoside (DDM) at purities ≥95% using a three-step process (Bumbak et al., 2018). ^1^H-^13^C HMQC spectra were collected for each construct in the apo, NT8-13 (agonist), SR142948A (antagonist) and ML314 (βArr-BAM) bound state; each newly observable resonance was then assigned to the respective residue (Figures 2A and S3-S6).

**Figure 2.**
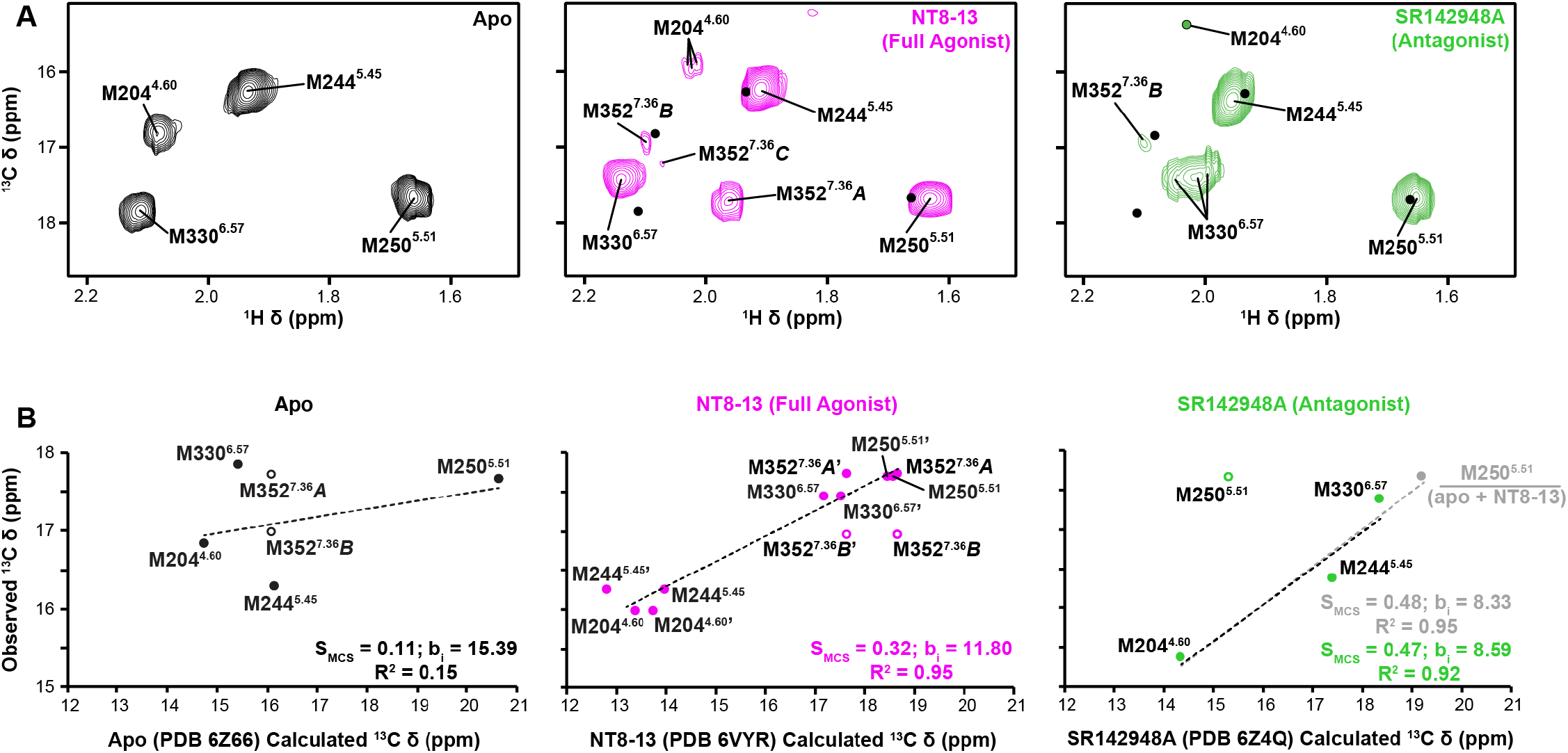
Ligand-induced chemical shift changes observed for ^13^CH_3_-methionine labelled enNTS_1_ΔM4. A) ^1^H-^13^C HMQC spectra of [^13^C^ε^H_3_-methionine]-enNTS_1_ΔM4 in the absence and presence of orthosteric ligands. All spectra were recorded at 600 MHz with protein concentrations of 66 μM. B) Linear correlation between DFT-calculated and experimental ^13^C chemical shift values. All observable resonances are included in the scatterplots. In spectra where a single methionine is assigned to multiple observable resonances, only the most populated (i.e. highest intensity) peak is fitted; the lower intensity peak is almost always an outlier (open circle). Linear regression of the DFT-calculated and observed ^13^C chemical shifts are generally well-correlated (0.86 ≤ R^2^ ≤ 0.95) apart from the apo-state (R^2^ = 0.15). The slope of the line (S_MCS_) can be interpreted as global side-chain order with values ranging from one (rigid) to zero (completely averaged). For the NT8-13-state, calculated chemical shifts from chain B of the X-ray structure are indicated with a prime. Because the M250^5.51^ sidechain in the SR142948A crystal structure (PDB 6Z4Q) was orientated drastically different from all other published crystal structures we fitted the linear regression two ways: (1; black line) excluding M250^5.51^, and (2; gray line) including the average of DFT-calculated chemical shifts for M250^5.51^ from apo and NT8-13-bound crystal structures.

### Ligands modulate local methionine sidechain dynamics

All NMR spectra were collected with identical acquisition parameters at 65 μM [^13^C^ε^H_3_-methionine]-enNTS_1_ΔM4; thus, both methionine methyl chemical shift values and signal intensities can be directly compared to monitor the effect of ligands. Intensity decreases are usually caused by line broadening and may reflect changes in the transverse relaxation rate (R_2_) and/or exchange broadening. R_2_ relaxation results from physical properties of the methyl group on the pico-nanosecond timescale, while exchange broadening reflects conformational interconversion (i.e. the methyl group sensing different chemical environments) in the micro-millisecond timescale. Throughout the subsequent results subsections, empirically observed changes in intensity (Figure S7) are either analyzed individually between two situations (e.g. ligand-free and ligand-bound) or pair-wise, by observing how intensities of two methionine signals respond to multiple changes in their chemical environments (e.g. apo and multiple ligands). For example, signal intensities for both M244^5.45^ and M250^5.51^ decrease (19% and 32%, respectively) upon addition of NT8-13 suggesting that the agonist induces microsecond-millisecond motions near the PIF motif (Figure S7). In contrast, SR142948A leaves the M250^5.51^ intensity unchanged from the apo-state while reducing the basal motions of M244^5.45^ and M330^6.57^ as reflected by peak intensity increases of 14% and 45%, respectively (Figure S7).

### Ligands tune global receptor flexibility

The numerous high-resolution NTS_1_ crystal structures available uniquely positioned us to interpret ligand-dependent methionine methyl carbon chemical shift values in terms of the S_MCS_ global order parameter. We focused our analyses on crystal structures of NTSR1-H4_X_, an evolved rNTS_1_ construct with 98% sequence identity to enNTS_1_ΔM4 across structured regions, in the apo (PBD 6Z66), NT8-13-bound (PDB 6VYR), and SR142948A-bound (PDB 6Z4Q) forms ^21^. All enNTS_1_ΔM4 methionine methyl groups are present in the crystallographic models, except for M352^7.36^ in the SR142948A-bound structure (PDB 6Z4Q). We confirmed that each structure was of sufficient refinement quality, specifically, possessing methionine S-C bond lengths between 1.77 and 1.80 Å ^20^. The eight residue substitutions unique to NTSR1-H4_X_ are all located beyond the 6 Å spheres around methionine methyl carbons and are unlikely to substantially contribute to the calculated chemical shift values.

Nearly all enNTS_1_ΔM4 NMR spectra contain multiple resonances assigned to a single methionine where the intensity of each resonance reflects its relative population at thermodynamic equilibrium (Figure 2A). The most intense peaks (i.e. most populated) show the best correlation with the DFT-calculated chemical shifts, confirming that the lowest energy conformers of the solution ensemble are frequently captured by X-ray structures (Figure 2B). The weaker alternative peaks reflect other well-populated conformations that are exchanging with the crystallographic conformer on the ms-s timescale (Figure 2B). Apo enNTS_1_ΔM4 possesses narrow, near random-coil chemical shift dispersion that produces a very low S_MCS_ = 0.11 (R^2^ = 0.15); this indicates a very flexible structure in which many other conformational states, beyond the one captured in the crystal structure, are present in solution. Addition of both orthosteric ligands dramatically increases the S_MCS_ indicating a substantial contraction of the receptor’s structural landscape (Figures 2B). The bulky SR142948A antagonist yields the most rigid enNTS_1_ΔM4 solution structure with a S_MCS_ = 0.47 when M250^5.51^ is excluded due to its weak electron density and unusual orientation (Figure 2B); if the predicted M250^5.51^ chemical shift is instead taken as the average of NT8-13 and apo structures, a similar S_MCS_ of 0.48 is obtained (Figure 2B). The NT8-13 full agonist produced slopes of 0.33 and 0.32 using the DFT-calculated chemical shift values from crystallographic chains A and B, respectively, which is comparable to the ligand binding domain of the glucocorticoid receptor ^20^. The high linear correlations between experimental and theoretical chemical shift values confirms i) that the low energy crystal structure conformation is largely populated in the solution ensemble, and ii) that the methionines, some of which are located 20-30 Å apart, sense common, collective motions across the receptor (Figure 2B). This provides confidence for interpreting the structural origin of ^13^C^ε^H_3_-methionine chemical shifts from the published crystal structures.

### Methionine probes of the orthosteric pocket inform on ligand association

Methionines 204^4.60^ and 208^4.64^ are located within the orthosteric pocket and are expected to be sensitive local probes for ligand binding (Figure 3D). In the apo-state, M204^4.60^ resonates in the same spectral region where the solvent-exposed methyl groups of M267^5.68^, M293^ICL3^ and M408^H8^ were observed prior to their removal (Figures 2A and S1). Inspection of the apo-state crystal structure clearly shows M204^4.60^ pointing into the empty binding pocket (Figure S8A). In the absence of direct interactions with other residues, the chemical shifts can be interpreted as indicating ∼40% χ^3^ trans conformation for apo-M204^4.60 29^; although it is generally recognized that the high flexibility of methionine methyl groups ^30^ and low energy barriers between rotamers ^29^ leaves structural interpretation of their ^13^C chemical shift values quite speculative. Regardless, the NMR chemical shifts, together with the strong resonance intensity (Figures 3A and S7) and the low global order parameter (Figure 2B), indicate apo-M204^4.60^ is highly mobile. Addition of NT8-13 or SR142948A induces sizeable upfield chemical shift perturbations (Figure 3A). The positions of the M204^4.60^ peak in the apo, NT8-13 and SR142948A bound forms are approximately aligned, despite the different nature of the ligands. Thus, it suggests the main effect of the ligands on M204^4.60^ chemical shifts is through the modulation of the contacts with other receptor residues, rather than the direct effect of the ligand on the methionine methyl group.

**Figure 3.**
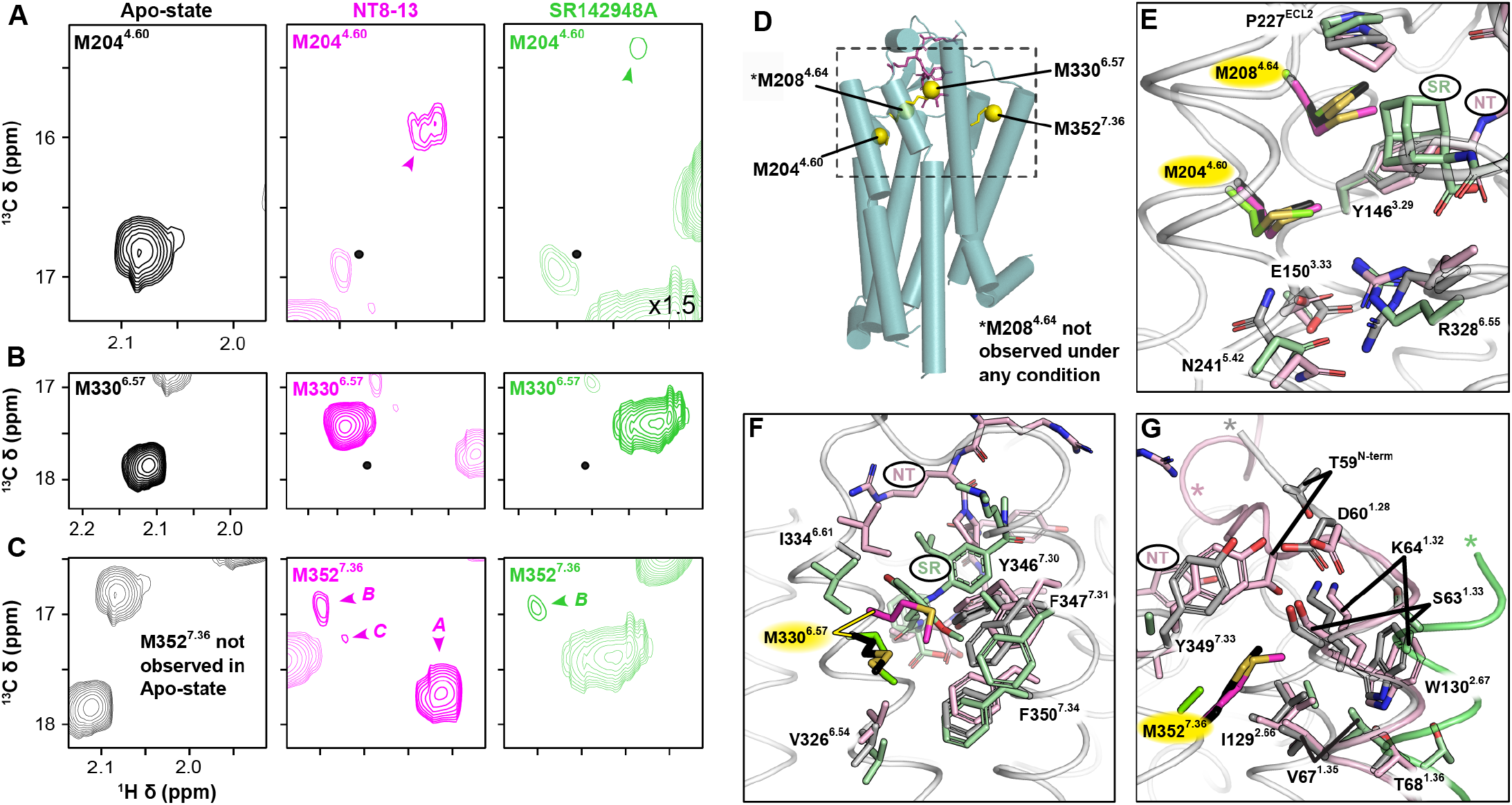
Ligand-dependent chemical shift perturbations in extracellular methionine residues. Extracted ^1^H-^13^C HMQC spectral regions of M204^4.60^ (A), M330^6.57^ (B), and M352^7.36^ (C) for apo (black), NT8-13 agonist-bound (magenta), and SR142948A antagonist-bound (green) enNTS_1_ΔM4; black dots in ligand-bound spectra denote the respective residue’s position in the apo-state. The resonances of other residues within the extracted region are drawn at 50% transparency. All spectra were recorded at 600 MHz with protein concentrations of 66 μM. D) Positions of extracellular methionine residues in thermostabilized rNTS_1_ (PDB 4BWB); M204^4.60^ resides within the orthosteric ligand binding site, M330^6.57^ at the extracellular tip of TM6 facing TM5, and M352^7.36^ at the extracellular tip of TM7 facing TM1. M208^4.46^ was not observed. E-G) Local environment of methionine residues observed in crystal structures of apo (methionine - black and surrounding residues - grey; PDB 6Z66), NT8-13-bound (methionine - magenta and surrounding residues - light pink; PDB 6YVR), and SR142948A-bound (methionine - green and surrounding residues - pale green; PDB 6Z4Q) NTS_1_-H4 ^21^. Portions of the ligands are labelled NT (NT8-13) and SR (SR142948A). In (E-F) the backbone trace is only shown for the apo-state (light grey). In (G) cartoons of TM1 and the N-terminus are illustrated for all structures. See Figure S8 for more detail.

M204^4.60^ lies adjacent to a hydrogen-bond network (E150^3.33^, T153^3.36^, Y154^3.37^, N241^5.42^, T242^5.43^ and R328^6.55^) that couples the orthosteric pocket to the connector region ^21^. Whereas the apo-state structure maintains a fully intact network involving all six amino acids (Figure S8A), binding of NT8-13 releases the E150^3.33^-T153^3.36^ and R328^6.55^-T242^5.43^ hydrogen-bonds leading to the formation of N241^5.42^-T153^3.36^ and N241^5.42^-R328^6.55^ interactions as well as a E150^3.33^-R328^6.55^ salt-bridge (Figures 3E and S8B). This reorganization positions the R328^6.55^ guanidinium group near the M204^4.60^ methyl (M204^4.60^C^ε^-R328^6.55^N^η2^ distance of 3.8 Å), which may explain the upfield chemical shift change in the ^13^C dimension ^20^. The upfield ^1^H chemical shift change may reflect the proximity of E150^3.33^O^ε1^ (4.1 Å vs. 5.2 Å in apo-state) as the distances and orientations of the other putative shielding residues, M208^4.64^ and Y146^3.29^, are nearly identical in the two structures. In the NT8-13-bound spectrum M204^4.60^ appears as three closely clustered peaks, which qualitatively reflects conformational exchange on the millisecond-second (i.e. slow) timescale. We hypothesize this results from fluctuations within the hydrogen-bond network induced by motions of the bound NT8-13 ligand ^31^ and/or remodeling of the connector region ^32^. SR142948A largely disrupts the network (Figure S8C), leaving only the N241^5.42^O^δ1^-R328^6.55^N^ε^ contact and a newly formed hydrogen bond between R328^6.55^N^η1^ and O5 of SR142948A. This sharply reduces the M204^4.60^C^ε^-R328^6.55^N^η2^ distance to 3.6 Å, which is consistent with its 0.6 ppm further upfield ^13^C chemical shift.

M208^4.64^ is the only resonance unobservable under all tested conditions. The M208^4.64^ methyl is stably sandwiched between the Y146^3.29^ and P227^ECL2^ sidechains, and then forms hydrophobic interactions with either the NT8-13 L13 C^δ^H_3_ group or the adamantane cage of SR142948A (Figures S8D-F). DFT calculations of these distinct chemical environments predict relative chemical shift values of -0.3, 0.29, and 0.84 ppm, respectively, from the NT8-13, apo, and SR142948A crystal structures. High microsecond to low millisecond exchange kinetics between these extreme chemical shifts would be expected to result in substantial exchange broadening; however, it’s also possible the signal is overlapped with DDM detergent resonances.

### The periphery of the orthosteric pocket undergoes slow timescale dynamics

M330^6.57^, positioned at the periphery of the orthosteric binding site between TM6 and TM7 (Figure 3), shows a complex response to the addition of the various ligands (Figure 3B). In the apo-state, M330^6.57^ manifests as a single, downfield resonance whose ^1^H chemical shift likely reflects subtle de-shielding from the F346^7.30^ and F350^7.34^ aromatic rings ^33^ although at 5.3 Å and 5.5 Å, respectively, their effect on the ^13^C chemical shift is minimal ^20^. The apo-M330^6.57^ carbon chemical shift reflects a near-equivalent 40:60 gauche:trans conformational equilibrium in the absence of significant neighbor-induced shielding effects ^29^. NT8-13 perturbs M330^6.57^ 0.03 ppm downfield in the ^1^H dimension and 0.4 ppm upfield in the ^13^C dimension (Figure 3B). This may reflect contraction of the orthosteric pocket and a concerted twisting of the M330^6.57^-bearing extracellular tip of TM6 that reduces M330^6.57^C^ε^-F346^7.30^C^ε2^, M330^6.57^C^ε^-F350^7.34^C^ε2/δ2^, and M330^6.57^C^ε^-Y347^7.31^C^δ1^ distances to 4.2, 3.5, and 5.5 Å, respectively (Figures S8G,H). SR142948A moves the M330^6.57 1^H frequency 0.1 ppm in the opposite direction (Figure 3B), consistent with the extracellular tips of TM6 and TM7 moving outwards to accommodate the bulky ligand (Figure 3F). Expansion of the orthosteric pocket increases the M330^6.57^C^ε^-F346^7.30^C^ε2^ separation to 5.8 Å while the F350^7.34^ sidechain re-orients to shield the M330^6.57^ methyl. This is accompanied by splitting of the resonance (Figure 2B) into at least three states qualitatively exchanging on the millisecond-second (i.e. slow) timescale. Van der Waals interactions with the dimethoxyphenyl group of SR148948A may further contribute to the complex behavior seen for M330^6.57^ (Figure S8I).

M352^7.36^ is situated at the extracellular tip of TM7 facing TM1 (Figure 3D) and is unobservable in the apo-state (Figure 3C). In the NT8-13 bound state, M352^7.36^ gives three peaks in slow exchange with intensities indicative of their relative population at thermodynamic equilibrium: states A (73.8%), B (13.9%), and C (12.3%). State B is the only observable resonance in the presence of SR142948A but with an absolute intensity 82% lower than with NT8-13 (Figures 3C and S7). NTS_1_ crystal structures reveal a substantial ligand-dependent rearrangement of the M352^7.36^ environment (Figures 3G and S8J-L). In the apo-structure, the M352^7.36^C^ε^ points toward the orthosteric binding site between TM7 and TM1 to interact with the aliphatic portion of the K64^1.32^ side chain (3.9 Å), S63^1.31^ (4.0 Å), I129^2.66^ (4.7 Å), V67^1.35^ (4.8 Å), and Y349^7.33^ (5.7 Å). D60^1.28^ plays a central role by engaging the S63^1.31^ hydroxyl and amide as well as the hydroxyl group of Y349^7.33^; this hydrogen-bond network loosely packs the extracellular tips of TM1 and TM7 with the receptor N-terminus. The N-terminus is largely unresolved, apart from N58 and T59, suggesting it does not stably associate with the orthosteric binding site in the apo-state.

In the NT8-13-bound structure, the receptor N-terminus extends over the orthosteric binding site to contact the ligand, extracellular loop (ECL)1, and ECL2. These extensive interactions pack the extracellular tips of TM1 and TM7 tightly around M352^7.36^, fully engaging the loose hydrogen bond network observed in the apo-state (Figure S8J,K). D60^1.28^, the first structured residue of TM1, pulls the N-terminus closer to the TM bundle through additional backbone and sidechain hydrogen bonds with I64^1.29^, S63^1.33^, K64^1.32^, and Y349^7.33^ (Figure S8K). A link between the extracellular tips of TM1 and TM2 is established via a T68^1.36^-W130^2.67^ hydrogen bond and hydrophobic interactions between the W130^2.67^ and K64^1.32^ sidechains. The K64^1.32^ sidechain forms an additional hydrogen bond with the carbonyl group of W130^2.67^, which is absent in the apo-state structure and suggests additional stabilization of the TM1-TM2 connection. In the SR142948A-bound crystal structure, the M352^7.36^ methyl group, the amino acid sidechains previously described within a 6 Å radius, the residues responsible for tethering the extracellular portion of TM1 to TM2/7, and the entire receptor N-terminus are all unresolved (Figure S8L) ^21^. Although SR142948A does not directly interact with TM1, this helix undergoes a substantial 3.8-5 Å outward translation as measured from the K64^1.32^ and S63^1.33^ C^α^ positions that expands the pocket and completely dissociates the N-terminus ^21^. This is also consistent with the deletion of rNTS_1_ N-terminal residues 45-60 reducing NT8-13 binding affinity >1000-fold but having no effect on SR48692 antagonist affinity _34_.

Taken together, we hypothesize that M352^7.36^ state A reflects the tightly packed TM1/TM2/TM7 interface and engaged receptor N-terminus that is only visible in the NT8-13-bound crystal structure whereas state B represents a dislocated N-terminus. State C likely reflects some intermediate combination of states A and B. This is supported by the high correlation between DFT-calculated chemical shifts and those observed for M352^7.36^ state A, but not state B (Figure 2B). The absence of M352^7.36^ in the apo-state spectrum likely signifies microsecond-millisecond timescale interconversion between an engaged and disengaged N-terminus. NT8-13 slows the overall exchange kinetics while stabilizing the engaged N-terminus whereas the neutral antagonist SR142948A prefers the disengaged position (Figure 3C).

### Methionine probes adjacent to the connector region report on ligand efficacy

The conserved PIF (P^5.50^/I^3.40^/F^6.44^) motif, a centrally located hydrophobic triad, connects the orthosteric pocket to the transducer binding site. Structures generally agree that agonist binding pulls the extracellular portion of TM5 inward, which forces I^3.40^ from between P^5.50^/F^6.44^ to permit outward rotation of TM6 ^9, 32, 35-36^. As such, ^13^C^ε^H_3_-methionine studies of other class A GPCRs have identified methionine probes on TM5 located below the P^5.50^ kink (M203^5.57^ in α_1A_AR ^37^, M223^5.54^ in β_1_AR ^38^, and M215^5.54^ in β_2_AR ^39^) as activation sensors that show efficacy dependent ligand-induced chemical shift changes. The PIF motif in enNTS_1_ΔM4 consists of P249^5.50^, A157^3.40^, and F317^6.44^ with two methionine probes in TM5, M244^5.45^ and M250^5.51^, located one turn prior to, and immediately following, the kink-inducing P249^5.50^ (Figure 4).

**Figure 4.**
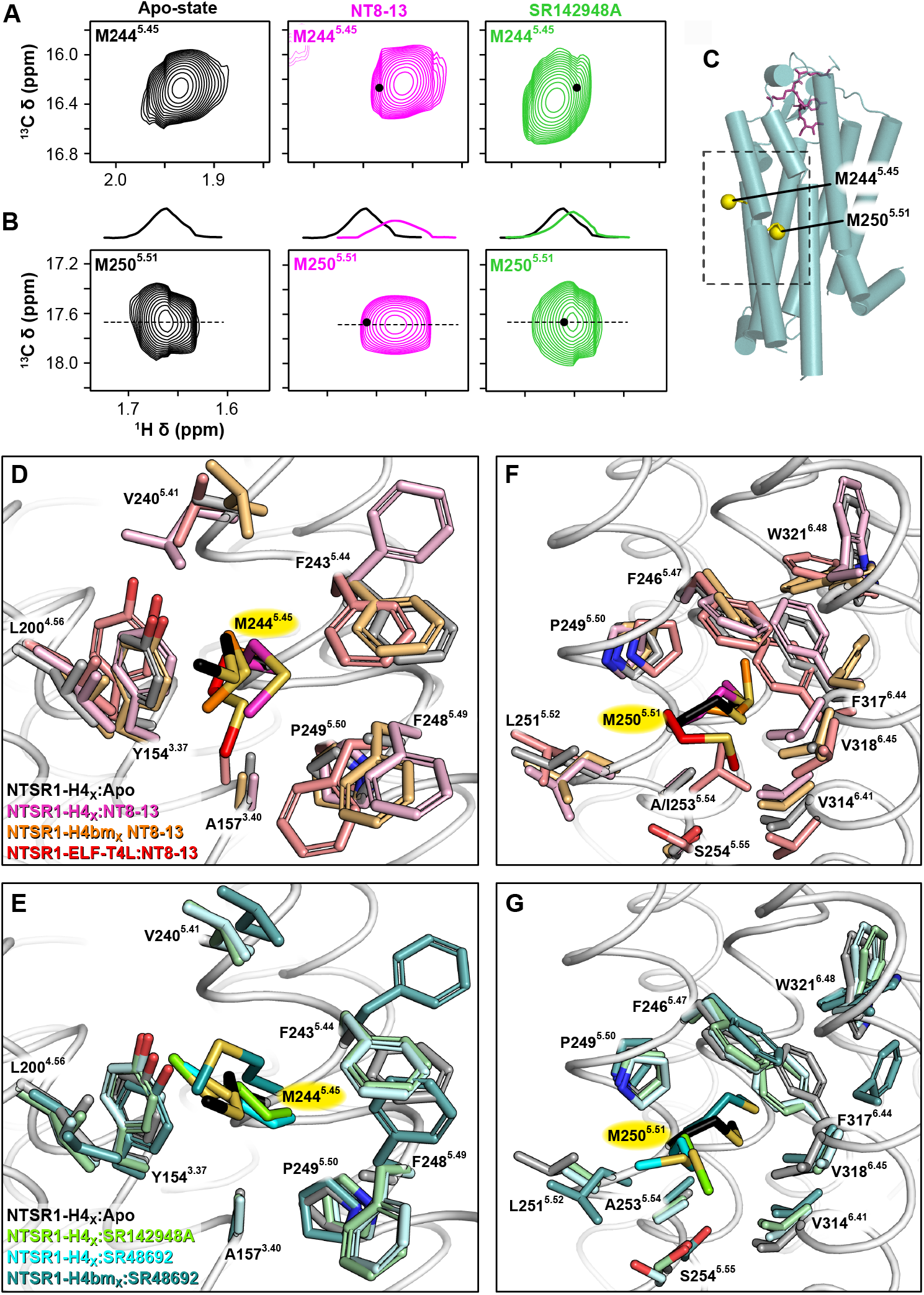
Ligand-dependent chemical shift perturbations for transmembrane domain methionine residues. Extracted ^1^H-^13^C HMQC spectral regions of M244^5.45^ (A) and M250^5.51^ (B) for apo (black), NT8-13 agonist-bound (magenta), and SR142948A antagonist-bound (green) enNTS_1_ΔM4; black dots in ligand-bound spectra denote the respective residue’s position in the apo-state. All spectra were recorded at 600 MHz with protein concentrations of 66 μM. C) Positions of transmembrane methionine residues in thermostabilized rNTS_1_ (PDB 4BWB); M244^5.45^ resides approximately one turn prior to the highly conserved P249^5.50^ and faces TM4 while M250^5.51^ is located immediately after P249^5.50^ and points directly into the PIF motif. D-G) Local environment of transmembrane methionine residues observed in crystal structures. Apo-state NTSR_1_-H4_X 21_ (methionine - black and surrounding residues - grey; PDB 6Z66). Only the backbone trace is shown for the apo-state (white cartoon). See Figures S9 and S10 for more detail. D & F) NT8-13 agonist-bound structures of NTSR_1_-H4_X_ (methionine - magenta and surrounding residues - light pink; PDB 6YVR ^21^), NTSR_1_-H4bm_X_ (methionine – orange and surrounding residues - light orange; PDB 6Z4V ^21^), and NTSR_1_-ELF-T4L (methionine - red and surrounding residues - salmon; PDB 4XEE ^76^). E & G) Antagonist-bound structures of NTSR_1_-H4_X_ in complex with SR142948A (methionine - green and surrounding residues - pale green; PDB 6Z4Q ^21^), NTSR_1_-H4_X_ in complex with SR48692 (methionine - cyan and surrounding residues - pale cyan; PDB 6ZIN ^21^), and NTSR_1_-H4bm_X_ in complex with SR48692 (methionine - teal and surrounding residues - light teal; PDB 6Z4S ^21^).

M244^5.45^ shows a clear linear response with NT8-13 inducing upfield ^1^H chemical shift perturbations compared to the apo-state, and SR142948A moving M244^5.45^ downfield (Figure 4A). The simplest explanation for this behavior is a two-state equilibrium (e.g. inactive-active) in the fast exchange NMR timescale that is modulated by ligand binding ^40^; this is consistent with SR142948A being an inverse agonist ^21, 41-42^ and the hypothesis that efficacy results from the ability of a ligand to stabilize a specific active state ^38-39, 43-45^. Structural analysis suggests that the M244^5.45^ chemical shift is primarily dominated by its proximity to Y154^3.37^. As mentioned earlier, the Y154^3.37^ sidechain is part of a hydrogen-bond network at the bottom of the orthosteric ligand binding pocket (Figure S8A-C). Except for two complexes, the majority of NTS_1_ structures suggest that the Y154^3.37^ hydroxyl group hydrogen bonds with orthosteric pocket residues, pulling it away from M244^5.45^, when the receptor is activated (Figure S9). This moves M244^5.45^ towards the shielding F248^5.49^ aromatic ring (Figures 4D and S9B-D). Whereas in the antagonist-bound structures, the M244^5.45^ sidechain is positioned closer to the edge of the Y154^3.37^ aromatic ring while F248^5.49^ moves out of the 6 Å sphere to produce an overall de-shielded environment relative to the apo-state (Figures 4E and S9E-G).

M250^5.51^ is positioned directly adjacent to P249^5.50^ of the PIF motif (Figures 4F,G and S10). It populates the most upfield ^1^H chemical shift observed in our study that, along with the ^13^C frequency, is relatively invariant regardless of ligand pharmacology (Figure 4B). Its unique resonance frequency is likely dominated by its proximity and orientation to the F317^6.44^ aromatic ring with subtle (de)shielding contributions from the F246^5.47^ and P249^5.50^ sidechains ^20^ (Figures 4F,G and S10). DFT calculations for the M250^5.51^ methyl group in the apo-state (PDB 6Z66) and NTSR1-H4_X_:NT8-13 (PDB 6YVR chains A and B) crystal structures predict that agonist binding induces a 0.034 ppm upfield shift in the ^1^H dimension, which is surprisingly consistent with the experimentally observed 0.03 ppm perturbation. SR142948A induces a subtle 0.01 ppm downfield ^1^H chemical shift perturbation in M250^5.51^, with respect to the apo-state spectrum, while the peak intensity remains virtually unchanged (Figures 4B and S7). Visual inspection of the NTSR1-H4_X_:SR142948A complex, and two structures with the chemically-related antagonist SR48692 ^21, 46^ (NTSR1-H4_X_:SR48692 and NTSR1-H4bm_X_:SR48692), reveals that the M250^5.51^ sidechain points away from the TM bundle with only V318^6.43^, S254^5.54^ and F246^5.47^ sidechains remaining within the 6 Å sphere (Figures 4G and S10E-G). Yet, the observed chemical shift is far more consistent with DFT calculations using the apo and NT8-13 structures with M250^5.51^ pointing towards the TM bundle (Figure 2B), which implies that the antagonist-bound crystal structures do not represent the full repertoire of NTS_1_ conformations present in solution.

### Methionine chemical shifts provide insight into BAM mechanism

To better understand the underlying mechanisms of ML314’s βArr-biased pharmacology ^27^, we collected ML314:enNTS_1_ΔM4 spectra in the presence and absence of NT8-13 (Figures 4 and S11A). Unfortunately, there are no published ML314:NTS_1_ complex structures to use for DFT calculations. If we instead substitute the NT8-13:NTSR1-H4_X_ crystal (PDB 6VYR) as a “rigid structure” reference then we obtain a S_MCS_ = 0.25 (Figure S11B). This suggests that the ML314 complex is more flexible than those of the orthosteric ligands, but more rigid than the apo receptor. Repeating this procedure with the ML314:NT8-13:enNTS_1_ΔM4 experimental chemical shifts produced a S_MCS_ of 0.34, which is slightly more rigid than NT8-13:enNTS_1_ΔM4 (Figure S11C). We can only speculate on ML314’s activation mechanisms without a published structure, however, our results may shed light on the binding interface itself. On addition of ML314, M330^6.57^ experiences an 0.03 ppm upfield ^1^H shift and 40% reduction in intensity compared to the apo-state (Figures 5B and S7D). Although the chemical shifts of M330^6.57^ in the apo and NT8-13 states are not the same, the peak shape and chemical shift perturbations induced upon the respective addition of ML314 are nearly identical. This consistent chemical shift change could result from a slight repositioning of the four nearby aromatic sidechains or as a direct result of ML314 binding. We hypothesize ML314’s molecular weight and hydrophobic character would target the receptor’s extracellular surface, perhaps interacting with the highly aromatic regions in the vicinity of TM6/7 or TM2/3.

**Figure 5.**
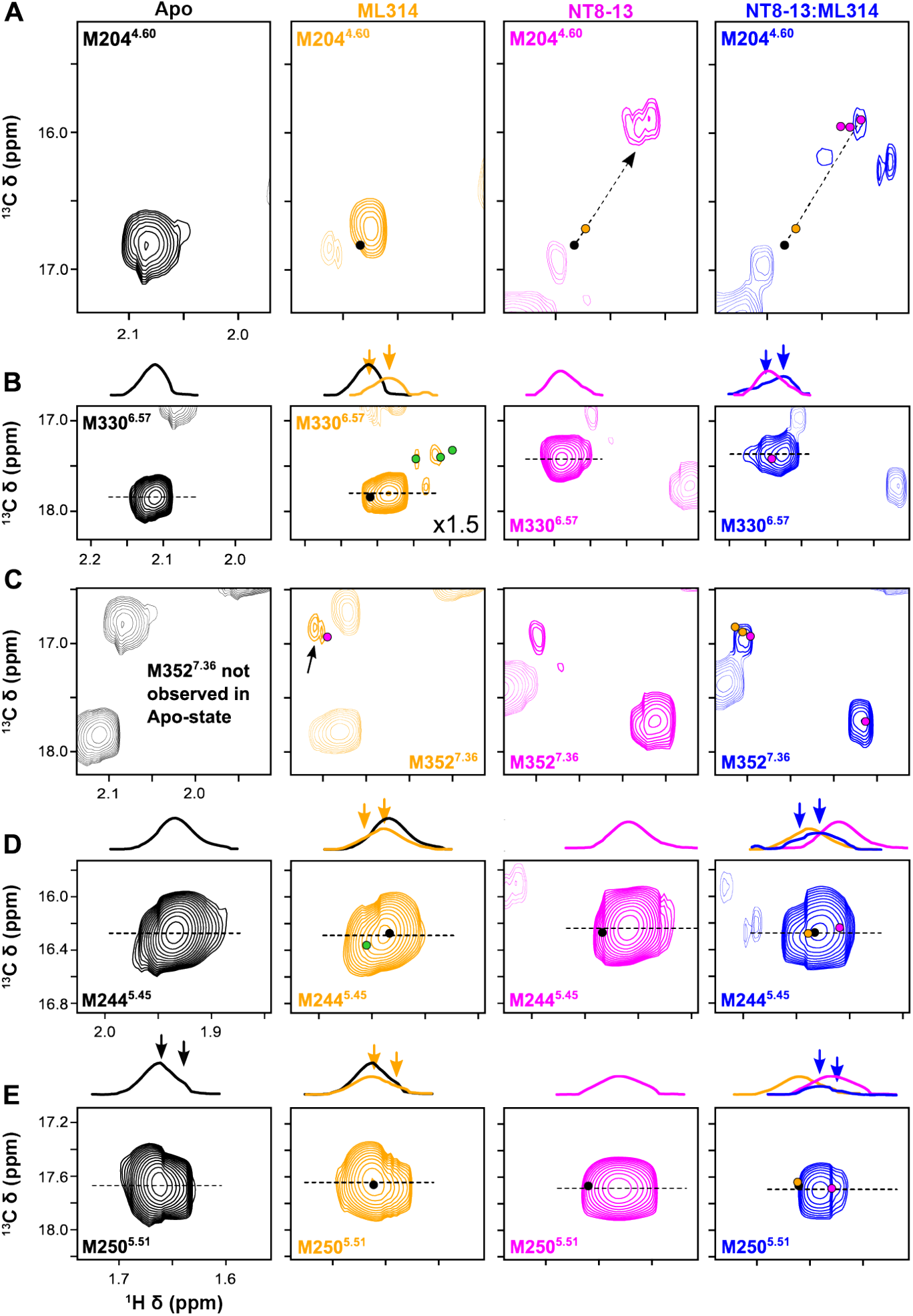
Methionine resonance perturbations near the connector region are suggestive of functional-selectivity. A-E) Extracted spectral regions of individual methionine residues for apo (black), M314 ago-BAM-bound (orange), NT8-13 agonist-bound (magenta), and ML314 and NT8-13 bound (blue) enNTS_1_ΔM4. The dots in ligand-bound spectra denote the respective residue’s position in the presence of alternative ligands (e.g. an orange dot marks the respective residue’s position in the ML314-bound spectrum). Resonances also observed in SR142948A antagonist-bound spectra are marked with green dots. All spectra were recorded at 600 MHz with protein concentrations of 66 μM.

The ML314 spectrum contains additional minor M330^6.57^ peaks. These signals appear at similar positions as the multiple peaks that are observed with SR142948A (Figures 3B and 5B; green dots). A similar antagonist-like effect is manifested in ML314’s selection of the M352^7.36^ B resonance (Figure 5C). On the co-addition of NT8-13 and ML314, M352^7.36^ state A and B (but not C) peaks are visible with reduced and increased intensities, respectively, compared to NT8-13 alone (Figures 7 and S11A). Crystal structures of receptors bound to pharmacologically-related positive allosteric modulators, such as the M2 muscarinic acetylcholine receptor complexed with LY2119620 ^47^, suggest a mechanism involving closure of the extracellular vestibule similar to transducer-bound conformations. However, SR142948A clearly expands the orthosteric pocket ^21^. This leads us to hypothesize that the similarity of SR142948A and ML314 complex spectra instead reflects detachment of the receptor N-terminus and local stabilization of extracellular TM1/6/7 (Figures 3C and 5C). ML314 may induce a conformation of the orthosteric binding pocket that allows a more favorable interaction with NT8-13 or even present a secondary NT8-13 binding site ^27^.

Conformational changes at M204^4.60^ near the bottom of the orthosteric pocket provide mechanistic hypotheses for BAM pharmacology ^27^. ML314 induces upfield shifts of M204^4.60^ in both dimensions and reduces the peak intensity by 54% (Figures 7A and S11A). These chemical shift changes appear along a linear trajectory defined by the apo- and NT8-13-bound states. The NT8-13:ML314:enNTS_1_ΔM4 spectrum closely resembles that of NT8-13 alone (Figure 11A); although the split M204^4.60^ peak is reduced to the most upfield resonance, perhaps mediating BAM-like outcomes, with a 45% reduction in signal intensity (Figure S7A). Three new resonances appear in the vicinity but cannot be assigned to M204^4.60^ based solely on their proximity (Figure 11A).

As the putative allosteric pipeline nears the connector region, resonances appear to exhibit signs of transducer functional selectivity. The M244^5.45^ chemical shifts do not change substantially from the apo-state, but ML314 reduces the peak intensity by 24% while giving rise to a nearby second peak (Figures 5D and S7B); again, at chemical shifts observed in the presence of SR142948A (Figure 5D; green dot). When both ligands are present, the peak doublet observed in the presence of ML314 is slightly exacerbated with corresponding increases in linewidths (Figure 5D). The ^1^H chemical shift value is centered halfway between those of either ligand individually; thus, in a sense ML314 antagonizes the chemical shift perturbation of NT8-13 (Figures 5D and S11A). Nearby M250^5.51^ responds similarly. ML314 has little effect on apo-state chemical shift but reduced the peak intensity by 26% (Figures 5E, S7C, and S11A). On addition of both NT8-13 and ML314, the M250^5.51 1^H chemical shift appears between the peak maxima of individual ligands while the peak intensity is dramatically reduced >60% (Figures 5E, S7C and S11A).

## DISCUSSION

Chashmniam *et al*. recently showed that the high intrinsic mobility of methionine methyl groups makes their ^13^C chemical shift sensitive to global sidechain motions ^20^. This observation is particularly powerful as the chemical shift is arguably the most accessible NMR-observable to quantitate. Analogous to their branched-chain counterparts, we hypothesize these methionine chemical shift-based global order parameters are, at least qualitatively, related to conformational entropy. Following this logic, molecular dynamics (MD) simulations and NMR relaxation data would further suggest that backbone motions play a limited role in the methionine chemical shift-based global order parameters ^14, 48^. Here, we tested whether methionine chemical shift-based order parameters (S_MCS_) can be used to discriminate global ligand-dependent GPCR dynamics.

The experimental ^13^C^ε^H_3_-methionine chemical shifts of enNTS_1_ΔM4 were compared to those calculated from high resolution crystal structures (Figures 2B and S11B,C). The striking linear correlation for NT8-13 and SR142948A complex chemical shifts establishes that i) the crystalized conformational states dominate the respective enNTS_1_ΔM4 solution ensembles, and ii) that well-distributed methionine residues are sensitive to collective GPCR motions on the sub-nanosecond timescale (Figure 6A). SR142948A dramatically reduces enNTS_1_ΔM4 global flexibility around a conformation incompatible with transducer complexation (Deluigi et al., 2021). Whereas, the agonist and BAM maintained relatively low global order parameters that are consistent with more frequent excursions to active-like conformers. If we assume that the other major entropic component, (de)solvation entropy ^49^, is not substantially different between orthosteric ligands it would suggest that antagonist binding is more enthalpically-driven. This would further indicate that NT8-13 remains relatively mobile upon complexation as previously suggested ^31^, and observed for the dynorphin/kappa opioid receptor ^50^. It is curious to note that the combined S_MCS_ of NT8-13 and ML314 is larger than either ligand individually, although not additive. We hypothesize this may reflect their underlying pharmacology where both ligands would be expected to stabilize a subset of partially overlapping fluctuations within the conformational ensemble. The narrow dispersion of apo enNTS_1_ΔM4 ^13^C chemical shifts were uncorrelated with the crystal structure suggesting a high degree of conformational averaging. Taken together, our results demonstrate that ligands selectively tune ps-ns dynamics, which suggests a role for dynamically-driven allostery in receptor activation (Figure 6A).

**Figure 6.**
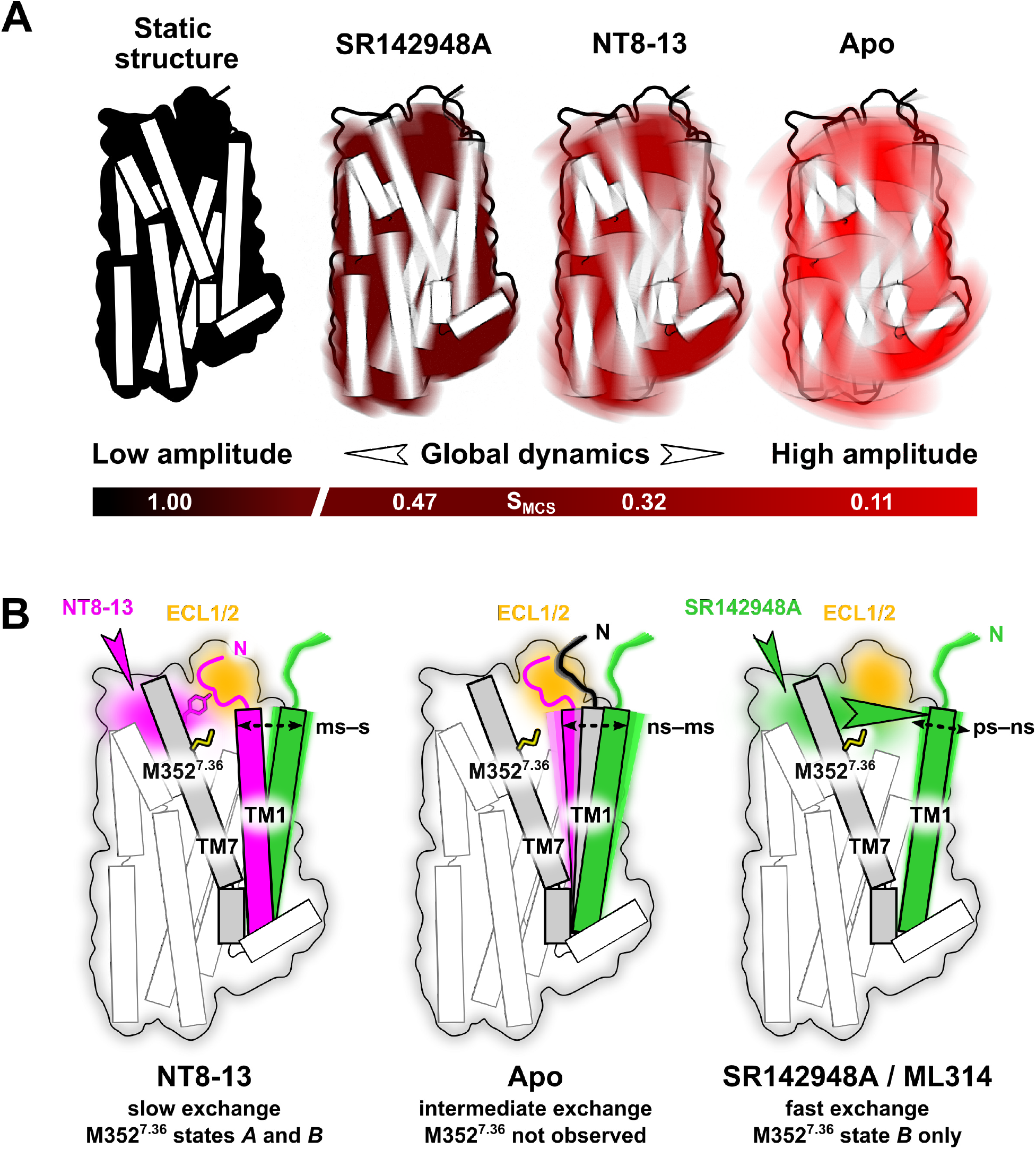
Illustration of ligand-mediated global and local GPCR dynamics. A) DFT calculations suggest that ligands differentially effect the amplitude of concerted enNTS_1_ motions as represented by the S_MCS_ global order parameter. B) The high linear correlation between DFT-calculated and experimentally observed chemical shifts lends confidence to spectral interpretation based upon the “rigid-limit” crystallographic structures.

The strong correlation between observed and DFT-calculated chemical shifts gave us confidence to interpret ligand-dependent perturbations in terms of their local electronic environment. From this, we can hypothesize on the effects of the yet biophysically-uncharacterized ML314 BAM. For example, M352^7.36^ senses a polar network that packs the TM1/7 extracellular tips against the N-terminus. Our solution observations agree with recent apo and antagonist-bound crystal structures that the N-terminus is primarily detached from the orthosteric binding site ^21^. While NT8-13 binding indeed stabilizes the NTS_1_ N-terminus with the extracellular vestibule in a lid-like manner, it continues to exchange slowly with a disengaged conformation reflecting previously observed flexibility in the orthosteric pocket (Figure 6B). SR142948A and ML314 both maintain the receptor with a detached N-terminus. Yet, ML314 leaves the orthosteric pocket empty while modestly rigidifying fast dynamics, which may provide a mechanism for potentiating NT8-13 efficacy ^26-27^. We identify M204^4.60^ of the orthosteric pocket as a sensor for subtle ligand-dependent rearrangements of a polar network that triggers the allosteric communication of extracellular ligand binding events to the transducer binding site. ML314 alone leads to an observable splitting and hence a reduction of M204^4.60^ exchange rate dynamics and may thus pre-select a NT8-13 competent conformation of the orthosteric binding site. M244^5.45^, located one turn above P^5.50^ of the conserved PIF motif (or connector region), exhibits clear efficacy-dependent chemical shift perturbations. While there is no comparable probe from previous ^13^C^ε^H_3_-methionine NMR studies, methionine residues after the P^5.50^ kink typically show strong correlations between chemical shift perturbations and ligand efficacy ^37-39, 51^; enNTS_1_ΔM4 M250^5.51^ does for NT8-13, but not SR142948A. While position 5.51 may be too close to the 5.50 hinge to sense the outward rotation of TM5, a similar behavior was observed for M2R M202^5.54 52^. Perhaps this reflects a general phenomenon in which receptors with shorter sidechains at position 3.40, V^3.40^ for M2R and A^3.40^ for NTS_1_, possess a less tightly packed (i.e. more dynamic) connector region compared to receptors with I^3.40^. Here, we demonstrate an approach for maximizing the information ^13^C^ε^H_3_-methionine probes provide on the thermodynamic, kinetic, and structural aspects of the free energy landscape.

A long-term goal of the GPCR biophysics community is to visualize the ligand binding and receptor activation process in the form of movies. While cryo-EM and crystallography have dutifully provided the actors, it’s up to NMR, fluorescence spectroscopy and MD to script their motions. The most-commonly applied NMR approach to characterize molecular motion is the model-free formalism developed by Lipari-Szabo ^53-54^. The resulting generalized order parameter serves as a dynamical proxy of conformational entropy ^14-16^, and thus a quantitative window into dynamically-driven allostery ^6, 12, 14, 55^. Accurate order parameter measurements require the nucleus of interest to be isolated from as many internuclear interactions (I.e. relaxation mechanisms) as possible ^56^. In large molecular weight systems, such as membrane proteins, this necessitates site-selective isotope labeling in highly-deuteration ^56-59^. Although this can be achieved in eukaryotic expression systems by elegant means ^60-62^, the ease with which isotopic labeling can be controlled in *E. coli* ^63^ uniquely poises amenable GPCRs for detailed characterization.

## Limitations of this study

We attribute the differential peak intensities to exchange-broadening motions on the microsecond-millisecond timescale (Figure S7). Although it is also possible that the ps-ns motions identified by DFT analysis could increase peak intensities through a longitudinal (T_1_) relaxation mechanism ^64^, we hypothesize that the SOFAST-HMQC experiments employed here greatly reduce that possibility ^65^. Nonetheless, future experiments will be required to quantitate the T_1_, T_2_, and generalized order parameters for each methionine methyl group. Our model system employed several strategies to minimize the challenges to solution NMR studies of GPCR complexes such as high molecular weights and inherent instability. To minimize line-broadening side effects of slowly tumbling systems, we employed DDM detergent micelles. Thermostabilized enNTS_1_ permits extended data acquisition times that would otherwise be impossible; it binds ligands with similar affinity to rNTS_1 66_, and couples directly to G protein and β-arrestin ^67^, although with reduced affinity (Figures 1 and S2). Future studies will explore reversion of thermostabilizing mutations to further recover signaling capabilities. Nonetheless, loss-of-function mutations are more common than gain-of-function phenotypes suggesting that the underlying mechanisms of enNTS_1_/transducer coupling represent native allosteric coupling.

## ACKNOWLEDGEMENTS

We would like to thank Prof. Andrew L. Lee (UNC) for fruitful discussions. The 14.1 T spectrometer at Indiana University used in this study was generously supported by the Indiana University Fund. The project was funded by: Indiana Precision Health Initiative (JJZ); NIH grants R00GM115814 (JJZ) and R35GM143054 (JJZ); KAKENHI 21H04791 (AI), 21H051130 (AI), JPJSBP120213501 (AI) from Japan Society for the Promotion of Science (JSPS); LEAP JP20gm0010004 (AI), BINDS JP20am0101095 (AI) from the Japan Agency for Medical Research and Development (AMED); FOREST Program JPMJFR215T (AI) and JST Moonshot Research and Development Program JPMJMS2023 (AI) from Japan Science and Technology Agency (JST); Daiichi Sankyo Foundation of Life Science (AI); Takeda Science Foundation (AI); Ono Medical Research Foundation (AI); and Uehara Memorial Foundation (AI); Agencia Estatal de Investigación, Spain grant PID2019-104914RB-I00 (MP, JCP).

## METHODS

### enNTS_1_ constructs and mutagenesis

The previously characterized functional variant enNTS_1_ (Bumbak et al., 2018) was available in an expression vector (termed pDS170) with an open reading frame encoding an N-terminal maltose-binding protein signal sequence (MBPss), followed by a 10x His tag, a maltose-binding protein (MBP), a NNNNNNNNNNG linker and a HRV 3C protease site (LEVLFQGP) which were linked via a BamHI restriction site (resulting in additional residues GS) to residue T42 of the receptor. C-terminally T416 of the receptor was linked via a NheI restriction site (resulting in additional residues AS) to an Avi-tag for in vivo biotinylation, a HRV 3C protease site, a GGSGGS linker and a monomeric ultra-stable green fluorescent protein (muGFP) ^68^.

The minimal methionine mutants enNTS_1_ΔM8A (M330^6.57^/M352^7.36^) and enNTS_1_ΔM8B (M204^4.60^/M208^4.64^) were selected ^69^ from a methionine mutant library obtained by DNA shuffling using the StEP ^70^. The DNA shuffling reaction was carried out using PrimeSTAR HS DNA polymerase (Takara, Mountain View, CA) and pDS170 plasmids carrying the parent templates enNTS_1_ΔM10 (containing no methionine residues, purchased form GenScript, Piscataway, NJ) and enNTS_1_ (containing the full set of 10 methionine residues) at a ratio of 5:1. The StEP library was amplified twice using Phusion HF DNA polymerase (NEB, Ipsiwtch, MA), separated on agarose gels and extracted using a Bioline Isolate II kit (Bioline, Taunton, MA). The amplified StEP library was then cloned into an expression vector containing a C-terminal mCherry instead of a muGFP fusion tag and transformed *E. coli* DH5α cells via electroporation at 2500 V using an Eppendorf 2150 electroporator (Eppendorf, Hauppauge, NY) and Biorad GenePulser cuvettes (Biorad, Hercules, CA). Electroporated cells were recovered in SOC medium for 1 h at 37 °C prior to further recovery in LB medium (containing 100 μg/ml ampicillin, 7% (w/v) sucrose, 1% (w/v) glucose) with shaking at 37ºC overnight. 15 mL of 2YT medium (containing 0.2% (w/v) glucose and 100 μg/μL ampicillin) were then inoculated with 7.5×10^8^ cells and grown to OD600 = 0.5 at 37 ºC with shaking. The temperature was then reduced to 20 ºC and StEP-NTS_1_ mutant library expression was induced with 250 μM IPTG. Expression was carried-out for approx. 16 h at 20 ºC with shaking. 12 mL of expression culture corresponding to 1.75×10^10^ cells, were centrifuged at 3000 rpm and 20 ºC (Thermo Multifuge), washed with 10 mL TKCl (50 mM Tris HCl, pH 7.4, and 150 mM KCl), resuspended in 10 mL TKCl and incubated for 2 h. The TKCl incubated cells were split in 2x 2 mL; to one tube (sort) 60 nM FAM-NT8-13 (0.86 μL) and to the other tube (competition) 60 nM FAM-NT8-13 (0.86 μL) and 10 μM NT8-13 (20 μL) were added and the mixtures were incubated in the dark for another 1 h at 20 ºC with shaking. The ligand-incubated cells were then centrifuged at 3000 rpm for 5 min, resuspended in 2 mL (sort) and 400 μL (competition) and filtered into FACS tubes. The cells incubated in the presence of competitor were used to calibrate the background for non-specific binding. From the other tube, 100,000 from the 8-10% most fluorescent cells were directly sorted into 5 mL LB medium containing 1% (w/v) glucose and recovered for 1 h at 37 ºC and shaking at 225 rpm. After recovery, 250 μL of the library were plated on an LB-agar plate containing 100 μg/mL ampicillin and 1% (w/v) glucose while 100 μg/mL ampicillin and incubated overnight at 37 ºC. Out of the >2000 colonies 24 were randomly picked and grown in 5 mL LB medium containing 100 μg/mL ampicillin and 1% (w/v) glucose overnight and the DNA was extracted for sequencing. enNTS_1_ΔM8A and enNTS_1_ΔM8B genes were extracted and cloned into the pDS170 expression vector containing the C-terminal muGFP gene as described before. To test expression and NT8-13 binding pDS170-enNTS_1_ΔM8A and pDS170-enNTS_1_ΔM8B were transformed into DH5α cells and each grown in 5 mL of 2YT containing 0.2% (w/v) glucose and 100 μg/μL ampicillin to OD600 = 0.5 at 37 ºC with shaking. The temperature was then reduced to 20 ºC and gene expression was induced with 250 μM IPTG. Expression was carried-out for approx. 16 h at 20 ºC with shaking. The next day, 2×10^8^ cells were centrifuged at 3000 rpm and 20 ºC (Thermo Multifuge) and resuspended in 2x 150 μL TKCl. To one of the aliquots 150 μL of 50 nM 5-TAMRA-NT8-13 in TKCl (sort) and to the other tube 150 μL of 50 nM 5-TAMRA-NT8-13 and 20 μM NT8-13 in TKCl (competition) were added and the mixtures incubated for 2 h at 20 ºC with shaking. The ligand-incubated cells were then centrifuged at 3000 rpm for 5 min, resuspended in 1 mL TKCl each and analysed using a LSR Fortessa X-20 FACS (BD Bioscience, Franklin Lakes, NJ). The mutant genes enNTS_1_ΔM7 (M204^4.60^/M330^6.57^/M352^7.36^), enNTS_1_ΔM6 (M204^4.60^/M208^4.64^/M330^6.57^/M352^7.36^), enNTS_1_ΔM5A (M204^4.60^/M208^4.64^/M244^5.45^/M330^6.57^/M352^7.36^), enNTS_1_ΔM5B (M204^4.60^/M208^4.64^/M244^5.45^ M250^5.51^/M330L/M352^7.36^) and enNTS_1_ΔM4 (M204^4.60^/M208^4.64^/M244^5.45^/M250^5.51^/M330^6.57^/M352^7.36^) were purchased from GenScript (Piscataway, NJ) and subcloned into the expression vector pDS170. All sub-cloning was done using the forward and reverse primers CATCATGGATCCACCTCTGAATCTGACACCGC and CATCATGCTAGCGGTAGAGAACGCGTGGTTAG, respectively and the restriction enzymes BamHI and NheI (NEB, Ipswitch, MA). The enNTS_1_ΔM5C (M204^4.60^/M208^4.64^/M244^5.45^ M250^5.51^/M330^6.57^/M352L) constructs were generated by site-directed mutagenesis based on enNTS_1_ΔM4 (M204^4.60^/M208^4.64^/M244^5.45^/M250^5.51^/M330^6.57^/M352^7.36^) using the Lightning site-directed mutagenesis kit (Agilent, Santa Clara, CA) and the forward and reverse primers CACTACTTCTACCTGCTGTCTAACGCGCTGG and CCAGCGCGTTAGACAGCAGGTAGAAGTAGTG, respectively.

### *E. coli* expression and purification of enNTS_1_ variants

The protocols for ^13^C^ε^H_3_-methionine labelled expression of enNTS_1_ variants used for all NMR experiments as well as unlabelled expression in rich media used for thermostability and binding assays have previously been described in depth ^66^. Expressions were usually carried out in batches of 3 L or 4 L and cell pellets were kept frozen at -80° C until further use. Purification of enNTS_1_, enNTS_1_ΔM1(M208V), enNTS_1_ΔM7 (M204^4.60^/M330^6.57^/M352^7.36^), enNTS_1_ΔM8A (M330^6.57^/M352^7.36^), enNTS_1_ΔM8B (M204^4.60^/M208^4.64^), enNTS_1_ΔM6 (M204^4.60^/M208^4.64^/M330^6.57^/M352^7.36^) and enNTS_1_ΔM5A (M204^4.60^/M208^4.64^/M244^5.45^/M330^6.57^/M352^7.36^) was carried out as described previously following a three-step purification protocol consisting of an immobilized metal affinity chromatography (IMAC) step for recombinant protein capture and exchange from n-decyl-β-D-maltopyranoside (DM) to DDM after solubilization, a reverse IMAC step to remove fusion proteins after cleavage with HRV 3C protease and a size exclusion chromatography (SEC) step ^66^. The variants enNTS_1_ΔM5B (M204^4.60^/M208^4.64^/M244^5.45^/M250^5.51^/M330L/M352^7.36^), enNTS_1_ΔM5C (M204^4.60^/M208^4.64^/M244^5.45^ M250^5.51^/M330^6.57^/M352L), enNTS_1_ΔM3 (M181^ICL2^, M204^4.60^/M208^4.64^/M244^5.45^/M250^5.51^/M330^6.57^/M352^7.36^), and the final variant enNTS_1_ΔM4 (M204^4.60^/M208^4.64^/M244^5.45^/M250^5.51^/M330^6.57^/M352^7.36^) were purified using a modified protocol. Elutions from the initial IMAC capture step were directly cleaved with His-tagged HRV 3C protease (produced in-house) prior to concentrating using an Amicon 30 kDa MWCO concentrator (Millipore) and dilution with ion exchange chromatography (IEX) loading buffer (20 mM HEPES pH 8.0, 10% Glycerol, 0.02% DDM) to obtain a combined NaCl/Imidazole/Na_2_SO_4_ concentration of less than 50 mM. The cleaved receptor solution was then loaded onto a 5 mL HiTrap SP HP column (GE Healthcare) using an Akta Start system (GE Healthcare) and washed with the same buffer until the signal remained stable. The column was then washed with four column volumes of IEX wash buffer (20 mM HEPES pH 7.4, 10% Glycerol, 63 mM NaCl, 0.02% DDM) after which a 1 mL Ni-NTA HisTrap column (GE Healthcare) was inserted after the HiTrap SP HP column and the system was washed with another 10 mL of IEX wash buffer containing 10 mM Imidazole. The cleaved receptor was the eluted with IEX elution buffer (20 mM HEPES pH 7.4, 10% Glycerol, 1 M NaCl, 0.03% DDM, 20 mM Imidazole) and the receptor containing fractions concentrated to approx. 400 uL for injection onto a S200 Increase SEC column (GE Healthcare) using a 500 μL loop and an Akta Pure System (GE Healthcare). The receptor containing fractions from SEC purification using SEC buffer (50 mM Potassium phosphate pH 7.4, 100 mM NaCl, 0.02% DDM) were then concentrated and buffer exchanged (for NMR experiments) using NMR buffer (50 mM Potassium phosphate pH 7.4, 100 mM NaCl in 100% D_2_O) to reduce the residual H_2_O concentration to <1%. Receptor samples were then aliquoted and stored at -80 ° C until further use. The modified purification protocol comprising the IEX step was found to yield a similar if not higher receptor purity compared to the original protocol containing a reverse IMAC step (Bumbak et al., 2019) as judged by SDS-Page. enNTS_1_ΔM4 used in NMR experiments retains a C-terminal Avi-tag (which was used for capture in ligand-binding and thermostability assays) and the amino acid sequence is: GPGSTSESDTAGPNSDLDVNTDIYSKVLVTAIYLALFVVGTVGNGVTLFTLARKKSLQSLQSRVDYYLGSLALSS LILLFALPVDVYNFIWVHHPWAFGDAGCKGYYFLREACTYATALNVVSLSVERYLAICHPFKAKTLLSRSRTKK FISAIWLASALLSLPMLFTMGLQNLSGDGTHPGGLVCTPIVDTATLRVVIQLNTFMSFLFPMLVASILNTVIARRLTVLVHQAAEQARVSTVGTHNGLEHSTFNVTIEPGRVQALRRGVLVLRAVVIAFVVCWLPYHVRRLMFVYISDEQWTTALFDFYHYFYMLSNALVYVSAAINPILYNLVSANFRQVFLSTLASLSPGWRHRRKKRPTFSRKPNSVSSNHAFSTASGLNDIFEAQKIEWHEGSGLEVLFQ

### NMR spectroscopy

NMR spectra were collected on 600 MHz Bruker Avance Neo spectrometers equipped with a triple resonance cryoprobes, except for the black spectra in Figure S1 and red spectra in Figures S3-6 which were recorded on a 800 MHz Bruker Avance II spectrometer, 2D ^1^H-^13^C SOFAST-HMQC spectra ^65^ were recorded with 25% non-uniform sampling (NUS) at 298 K with a ^1^H spectral width of 12 ppm (1024 data points in t_2_) and a ^13^C spectral width of 25 ppm (128 data points in t_1_), relaxation delays of 400 ms (800 MHz) and 450 ms (600 MHz), and 2048 scans per t1 data point resulting in acquisition times of 8 h (800 MHz) and 10 h (600 MHz) per spectrum. A 2.25 ms PC9 120 degree ^1^H pulse ^71^ was applied for excitation and a 1 ms r-SNOB shaped 180 degree ^1^H pulse ^72^ was used for refocusing. The ^13^C carrier frequency was positioned at 17 ppm, and the ^1^H at 4.7 ppm, while band selective ^1^H pulses were centered at 2 ppm (800 MHz) and 1.8 ppm (600 MHz). 1D ^1^H spectra were recorded at 298 K with a spectral width of 13.7 ppm (2048 data points) and a relaxation delay of 1 s, and 128 scans. Samples measured at 800 MHz were prepared to volumes of 290 μL in 5 mm Shigemi NMR tubes (Shigemi Inc., Allison Park, PA). Samples measured at 600 MHz were prepared to volumes of 160 μL in 3 mm tubes (Willmad). Samples measured at 600 MHz contained 20 μM DSS and 0.05% Na_2_N. Ligands were added to a final concentration of 500 μM. NT8-13 (5-10 mM) and SR142948A (20 mM) stock solutions were prepared in 100% D_2_O and ML314 (20 mM) in 100% DMSO-d6. Spectra measured at 600 MHz were referenced against internal DSS and spectra measured at 800 MHz were referenced against D_2_O. All spectra reconstructed with compressed sensing using qMDD ^73^ and processed using NMRPipe ^74^ where data were multiplied by cosinebells and zero-filled once in each dimension. Spectra were analyzed in Sparky (Goddard, T.D. and Kneller, D.G., University of California, San Francisco). All spectra in this study are reproduced together in Figure S12.

### TGFα shedding assay

TGFα shedding assay was performed as described previously ^24^. HEK293A cells (Thermo Fisher Scientific) were seeded in a 6-well culture plate (Greiner Bio-One) at a concentration of 2×10^5^ cells/ml (2 mL per well hereafter) in DMEM (Nissui Pharmaceutical) supplemented with 10% (v/v) FBS (Gibco), glutamine, penicillin, and streptomycin, one day before transfection. The transfection solution was prepared by combining 5 μL of 1 mg/ml polyethylenimine Max solution (Polysciences) and a plasmid mixture consisting of 200 ng ssHA-FLAG-NTS_1_ and 500 ng alkaline phosphatase (AP)-tagged TGF-α (AP-TGFα; human codon-optimized). One day after incubation, the transfected cells were harvested by trypsinization, neutralized with DMEM containing 10% (v/v) FCS and penicillin–streptomycin, washed once with Hank’s Balanced Salt Solution (HBSS) containing 5 mM HEPES (pH 7.4), and resuspended in 6 ml of the HEPES-containing HBSS. The cell suspension was seeded into a 96-well plate at a volume of 80 μL (per well hereafter) and incubated for 30 minutes in a CO_2_ incubator. A test ligand (ML314; diluted in 0.01% (w/v) BSA and 5 mM HEPES-containing HBSS (assay buffer) at 10x concentration) or vehicle was added at a volume of 10 μl. After 5 min, a test agonist (NT8-13; serially diluted in assay buffer at 10x concentration) was added and the plate was incubated for 1 h. After centrifugation, conditioned media (80 μL) was transferred to an empty 96-well plate. AP reaction solution (10 mM p-nitrophenylphosphate (*p*-NPP), 120 mM Tris–HCl (pH 9.5), 40 mM NaCl, 10 mM MgCl_2_) was dispensed into the cell culture plates and plates containing conditioned media (80 μl). Absorbance at 405 nm was measured before and after a 1 h or 2 h incubation at room temperature using a microplate reader (SpectraMax 340 PC384; Molecular Devices). Ligand-induced AP-TGF-α release was calculated as described previously.^26^ Unless otherwise noted, vehicle-treated AP-TGF-α release signal was set as a baseline. Using Prism 8 software (GraphPad Prism), AP-TGF-α release signals were fitted with a four-parameter sigmoidal concentration-response curve.

### NanoBiT-based βArr1 assay

NanoBiT-based βArr1 assay was performed as described previously ^75^. HEK293A cells were seeded in a 6 cm culture dish (Greiner Bio-One) at a concentration of 2×10^5^ cells/mL (4 mL per dish) in the FBS-supplemented DMEM. Plasmid transfection was performed by combining 10 μL of the polyethylenimine Max solution and a plasmid mixture consisting of 1 μg ssHA-FLAG-GPCR-SmBiT and 200 ng LgBiT-βArr1 in 400 μL of Opti-MEM. After an incubation for one day, the transfected cells were harvested with 0.5 mM EDTA-containing Dulbecco’s PBS (D-PBS), centrifuged, and suspended in 4 mL of HBSS containing 0.01% (w/v) BSA and 5 mM HEPES (pH 7.4) (assay buffer). The cell suspension was dispensed in a white 96-well plate (Greiner Bio-One) at a volume of 70 μL per well and loaded with 20 μL of 50 μM coelenterazine (Carbosynth), diluted in the assay buffer. After 2 h incubation at room temperature, the plate was measured for its baseline luminescence (SpectraMax L, 2PMT model, Molecular Devices). Thereafter, a test ligand (ML314; diluted in assay buffer at 10x concentration) was added at a volume of 10 μL and the plate was incubated for 15 min at room temperature. After a second measurement of luminescence, a test agonist (NT8-13; serially diluted in assay buffer at 6x concentration) were added at a volume of 20 μL and the plate was immediately read as a kinetics mode for 10 min. Luminescence counts recorded from 5 min to 10 min after the agonist addition were averaged and normalized to the initial counts. The fold-change signals were further normalized to the vehicle-treated signal and were plotted as a βArr1 recruitment response. Using the Prism 8 software, the βArr1 recruitment signals were fitted to a four-parameter sigmoidal concentration-response curve.

### Flow cytometry analysis

Plasmid transfection into HEK293A cells were performed as described in the TGFα shedding assay section. One-day after transfection, the cells were collected by adding 200 μL of 0.53 mM EDTA-containing D-PBS, followed by 200 μL of 5 mM HEPES (pH 7.4)-containing HBSS. The cell suspension was transferred to a 96-well V-bottom plate in duplicate, blocked with 2% (v/v) goat serum- and 2 mM EDTA-containing D-PBS (blocking buffer; 100 μL per well hereafter) and fluorescently labeled with the anti-FLAG-epitope tag monoclonal antibody (Clone 1E6, FujiFilm Wako Pure Chemicals; 10 μg per ml in the blocking buffer; 25 μL) and a goat anti-mouse IgG secondary antibody conjugated with Alexa Fluor 488 (Thermo Fisher Scientific, 10 μg per mL diluted in the blocking buffer; 25 μL). Live cells were gated with a forward scatter (FS-Peak-Lin) cutoff at the 390 setting, with a gain value of 1.7 and fluorescent signal derived from Alexa Fluor 488 was recorded in the FL1 channel.

## Supplemental Information

**Supplemental Figure S1.**
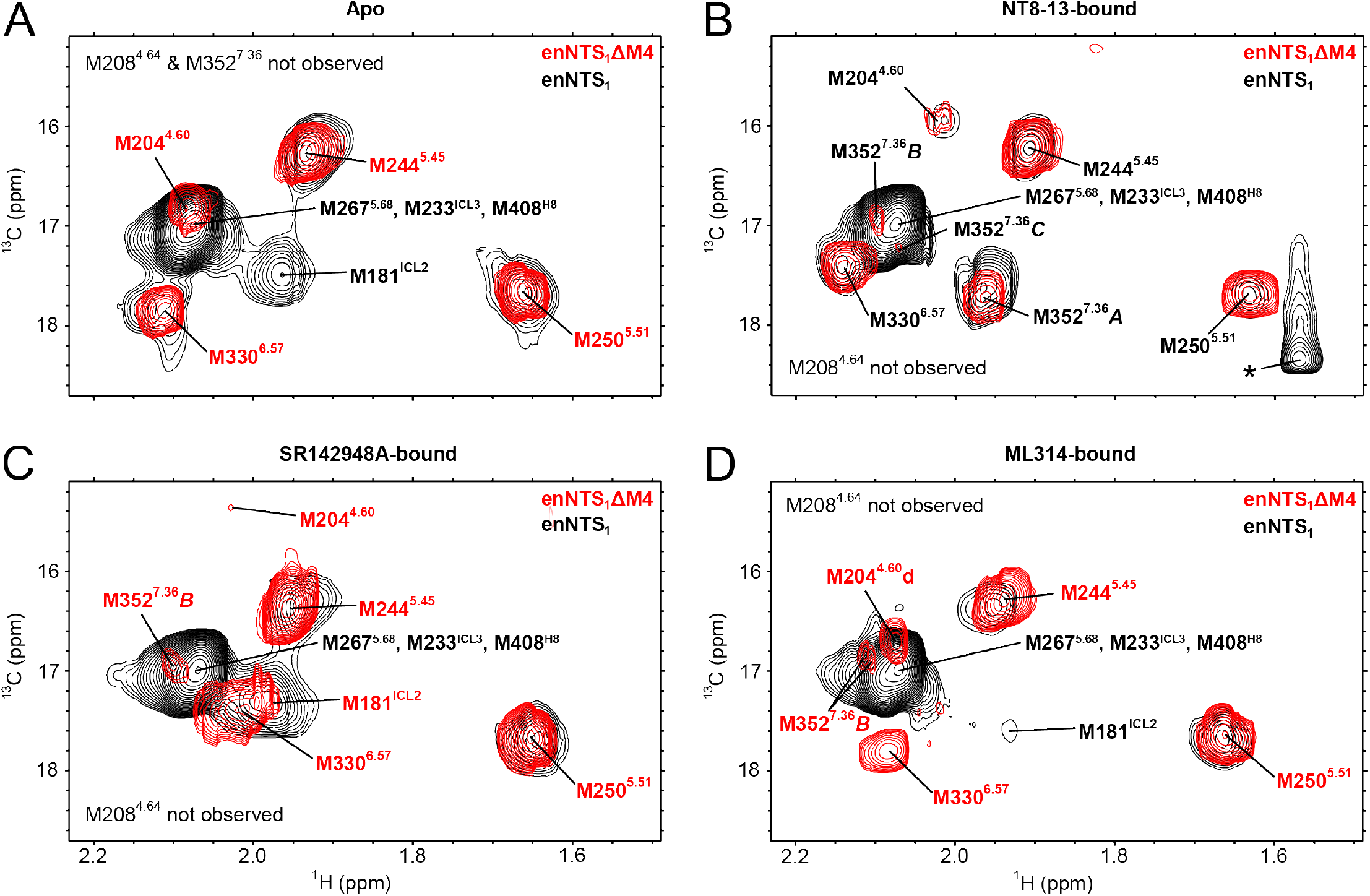
Site-selective methionine mutagenesis has little to no effect on ligand-dependent chemical shifts. Overlay of ^13^C^ε^H_3_-methionine-labeled enNTS_1_ΔM4 (red) and enNTS_1_ (black) ^1^H-^13^C HMQC spectra for apo (A), NT8-13-bound (B), SR142948A-bound (C), and ML314-bound (D) conditions. M267^5.68^, M293^ICL3^, and M408^H8^ of enNTS_1_ are likely solvent exposed and form an intense cluster of resonances overlapping with M204^4.60^ in Apo and ML314-bound states, M352^7.36^B in NT8-13-bound, SR142948-bound, and ML314-bound states, M352^7.36^B in the NT8-13-bound state, and M330^6.57^ in NT8-13-bound and SR142948A-bound states. M181^ICL2^ of enNTS1 overlaps with M330^6.57^ in the SR142948A-bound state and with M352^7.36^A in NT8-13-bound state. M208^4.64^ was not observed in any spectra while M352^7.36^ was not observed in Apo-state spectra. The asterisk in panel B marks a contaminant peak which was observed in all assignment spectra containing NT8-13 (also see Supplemental Figure S4). The most notable differences are observed for the ML314 spectra (D) where a) the enNTS_1_ but not the enNTS_1_ΔM4 spectrum shows a complete disappearance of the main M330^6.57^ resonance and b) M181^ICL2^ of enNTS_1_ significantly broadens while moving upfield in the ^1^H dimension. Possible reasons for these differences are discussed in the ML314 section of the main manuscript. enNTS_1_ΔM4 spectra (red) were recorded at 600 MHz, in 3 mm thin wall precision NMR tubes (Wilmad), with protein concentrations of 66 μM and enNTS_1_ spectra (black) were recorded at 800 MHz, in 5 mm Shigemi tubes, with protein concentrations of 28 μM.

**Supplemental Figure S2.**
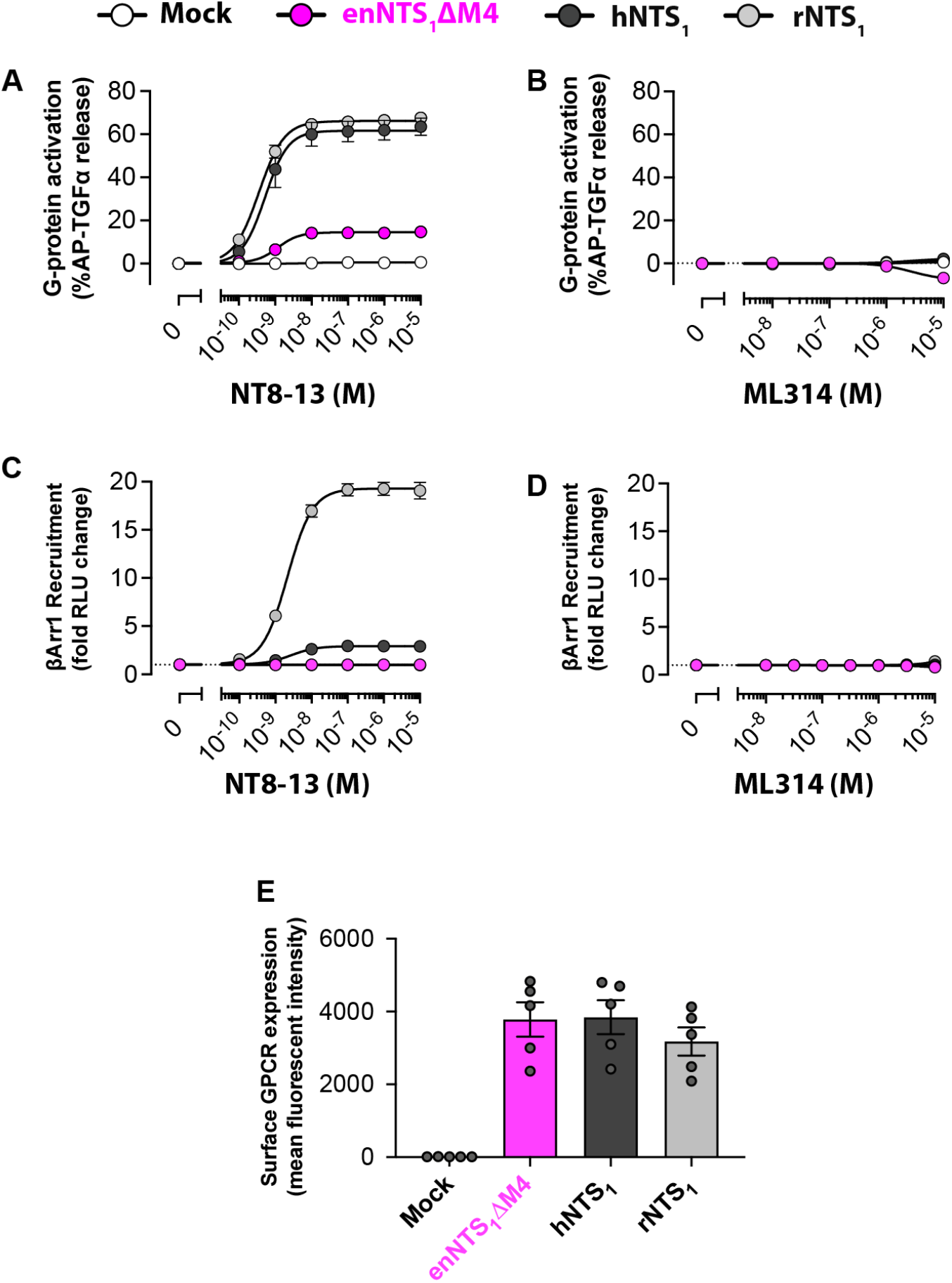
Cell-based G protein activation and βArr1 recruitment. A) NT8-13 dependent and B) ML314 dependent G protein activation of mock transfection (open), enNTS_1_ΔM4 (magenta), rNTS_1_ (gray), and hNTS_1_ (black) in a TGFα shedding assay using HEK293A cells ^1^. Error bars represent SEM from three independent experiments. C) NT8-13-dependent and D) ML314-dependent βArr1 recruitment to mock transfection (open), enNTS_1_ΔM4 (magenta), rNTS_1_ (gray), and hNTS_1_ (black) in a NanoBiT-based assay using HEK293A cells ^2^. Error bars represent SEM from three independent experiments. Note that enNTS_1_ΔM4 was unresponsive in the β-arrestin assay and thus the data points overlap with the mock responses. Also note that for many samples in A-D, the size of error bar is smaller than the symbol. E) Surface expression levels for each receptor were measured using flow cytometry. Bars and error bars represent mean and SEM from five independent experiments (dots).

**Supplemental Figure S3.**
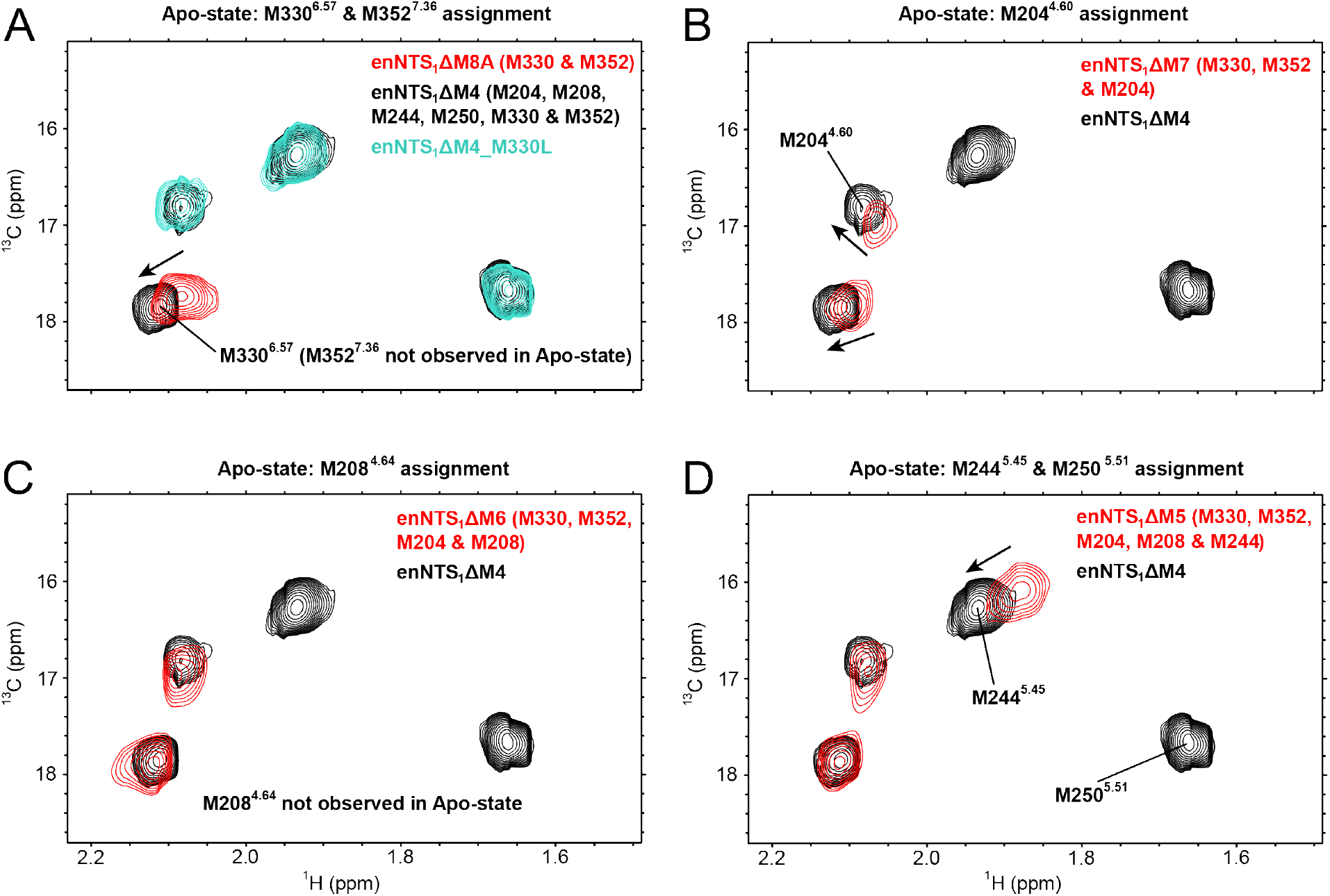
Assignment of apo enNTS_1_ΔM4 ^13^C^ε^H_3_-methionine resonances. ^1^H-^13^C HMQC spectra of ^13^C^ε^H_3_-methionine-labeled enNTS_1_ΔM4 (black) and its variants (red or turquoise); peaks that appear (disappear) with each amino acid substitution were assigned to mutated residues as labelled. All spectra were collected in the apo state. enNTS_1_ΔM4 (black) and enNTS_1_ΔM4_M330L (turquoise) spectra were recorded at 600 MHz, in a 3 mm thin wall precision NMR tube (Wilmad), with protein concentrations of 66 μM and 73 μM, respectively. enNTS_1_ΔM8A, ΔM7, ΔM6 and ΔM5 spectra (red) were recorded at 800 MHz, in 5 mm Shigemi tubes, with protein concentrations of 20 μM, 20 μM, 21 μM, and 24 μM, respectively.

**Supplemental Figure S4.**
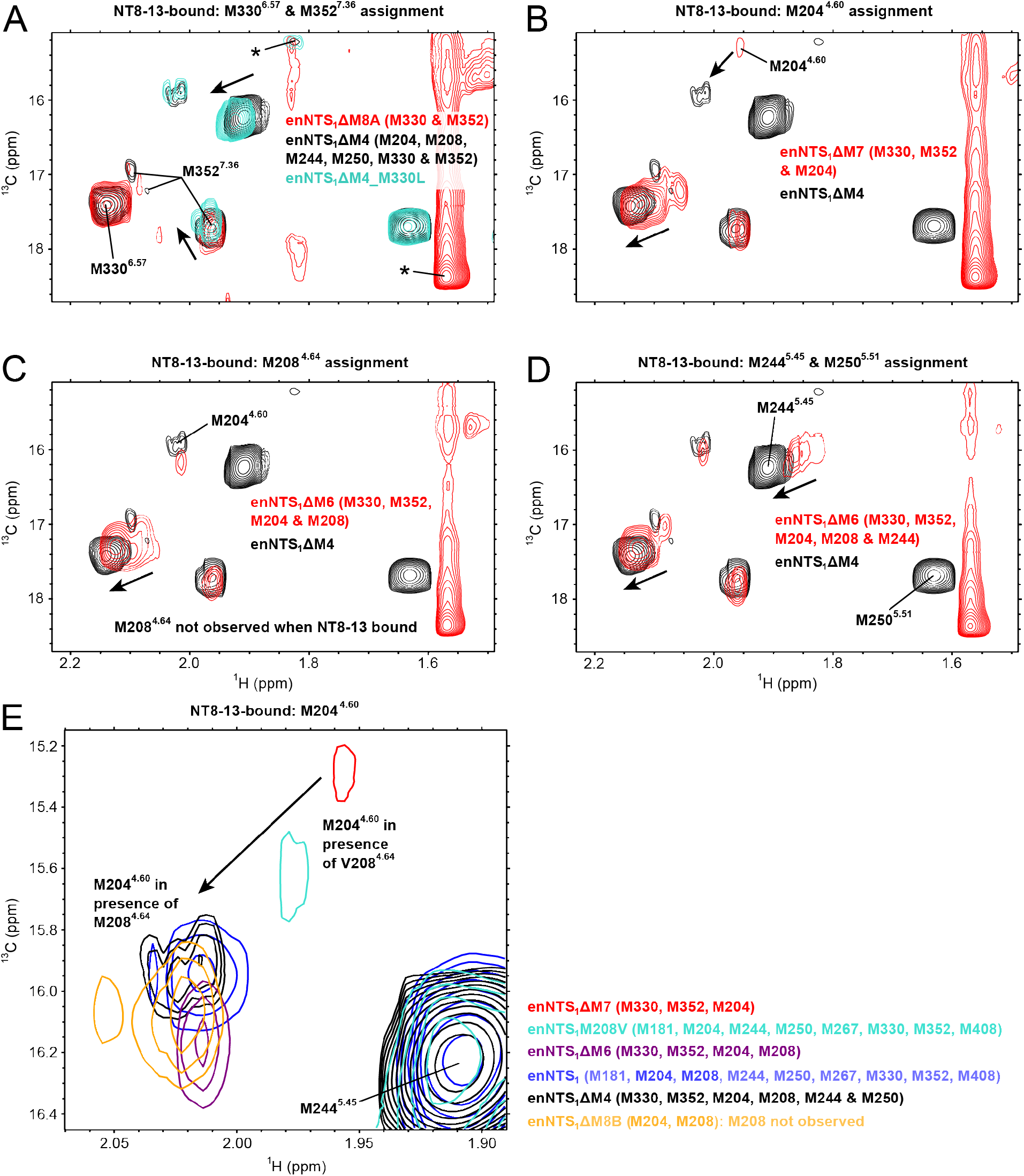
Assignment of NT8-13-bound enNTS_1_ΔM4 ^13^C^ε^H_3_-methionine resonances. ^1^H-^13^C HMQC spectra of ^13^C^ε^H_3_-methionine-labeled enNTS_1_ΔM4 (black) and its variants (colors); peaks that appear (disappear) with each amino acid substitution were assigned to mutated residues as labelled. All spectra were collected in the apo state. enNTS_1_ΔM4 (black) and enNTS_1_ΔM4_M330L (turquoise) spectra were recorded at 600 MHz, in a 3 mm thin wall precision NMR tube (Wilmad), with protein concentrations of 66 μM and 73 μM, respectively. enNTS_1_ΔM8A, ΔM7, ΔM6, ΔM5 spectra (red), and enNTS_1_ΔM8A (orange) were recorded at 800 MHz, in 5 mm Shigemi tubes, with protein concentrations of 20 μM, 20 μM, 21 μM, 24 μM, and 36 μM respectively.

**Supplemental Figure S5.**
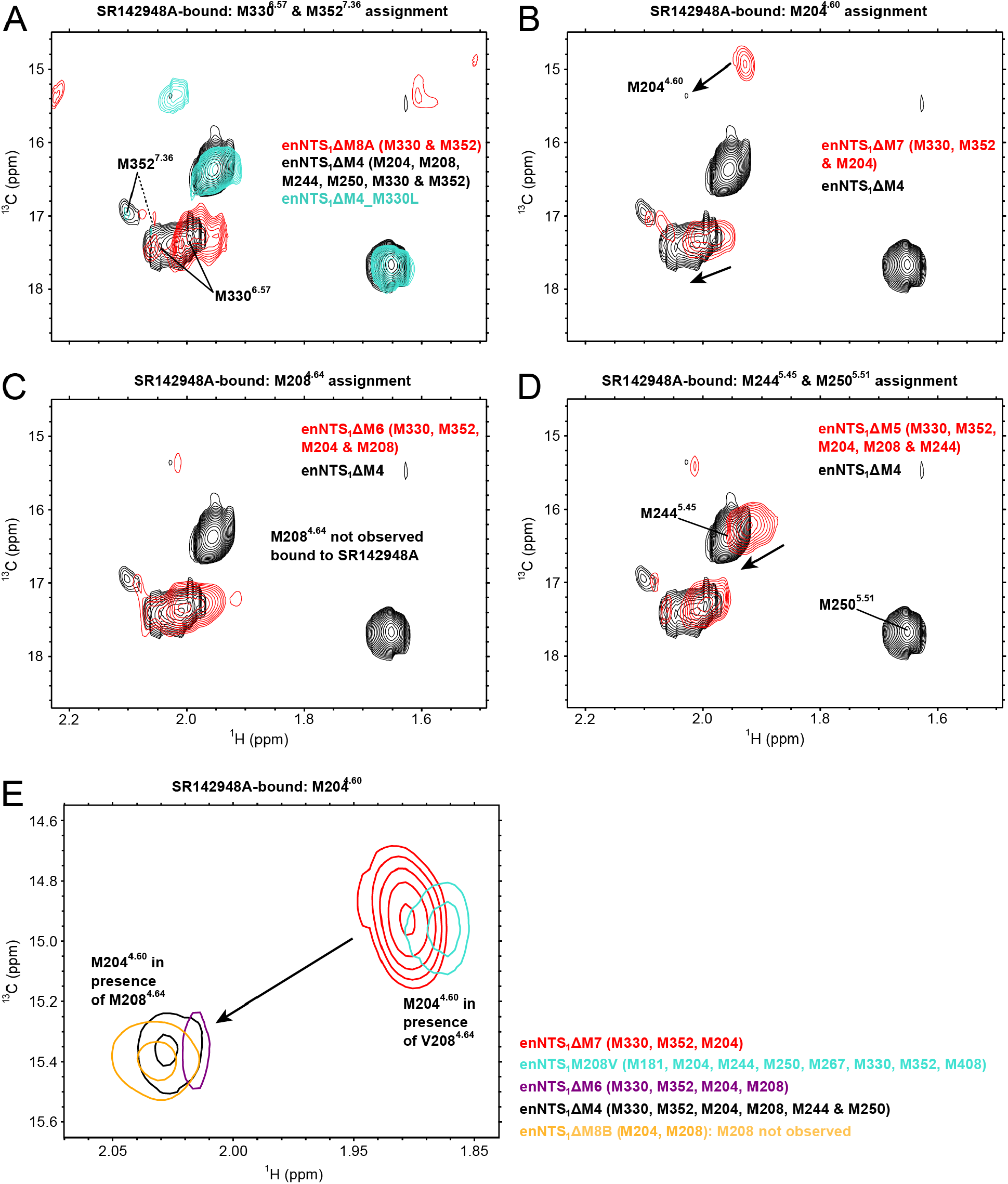
Assignment of SR142948A-bound enNTS_1_ΔM4 ^13^C^ε^H_3_-methionine resonances. ^1^H-^13^C HMQC spectra of ^13^C^ε^H_3_-methionine-labeled enNTS_1_ΔM4 (black) and its variants (colors); peaks that appear (disappear) with each amino acid substitution were assigned to mutated residues as labelled. All spectra were collected in the apo state. enNTS_1_ΔM4 (black) and enNTS_1_ΔM4_M330L (turquoise) spectra were recorded at 600 MHz, in a 3 mm thin wall precision NMR tube (Wilmad), with protein concentrations of 66 μM and 73 μM, respectively. enNTS_1_ΔM8A, ΔM7, ΔM6, ΔM5 spectra (red), and enNTS_1_ΔM8A (orange) were recorded at 800 MHz, in 5 mm Shigemi tubes, with protein concentrations of 20 μM, 20 μM, 21 μM, 24 μM, and 36 μM respectively.

**Supplemental Figure S6.**
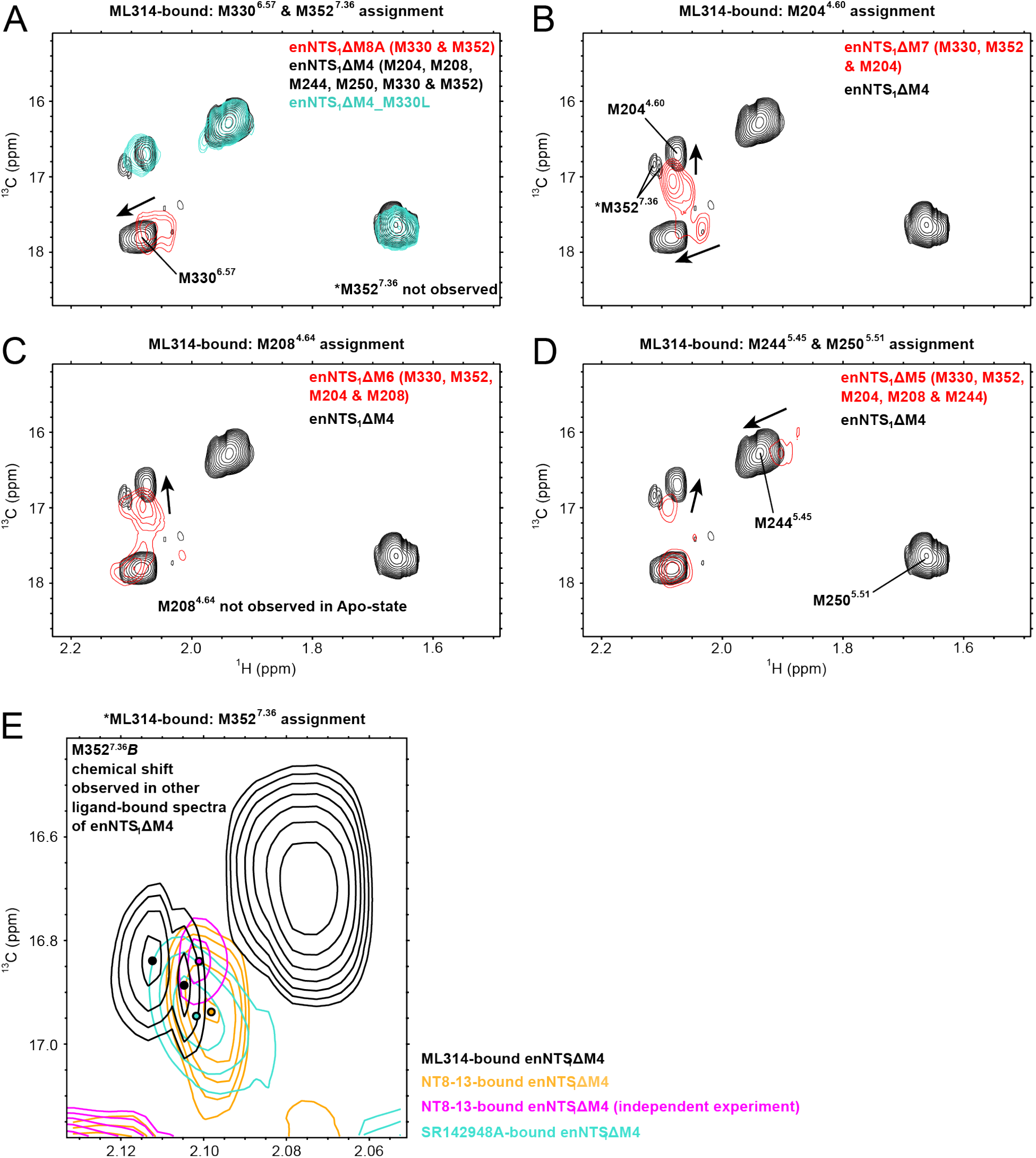
Assignment of ML314-bound enNTS_1_ΔM4 ^13^C^ε^H_3_-methionine resonances. ^1^H-^13^C HMQC spectra of ^13^C^ε^H_3_-methionine-labeled enNTS_1_ΔM4 (black) and its variants (red or turquoise); peaks that appear (disappear) with each amino acid substitution were assigned to mutated residues as labelled. All spectra were collected in the apo state. enNTS_1_ΔM4 (black) and enNTS_1_ΔM4_M330L (turquoise) spectra were recorded at 600 MHz, in a 3 mm thin wall precision NMR tube (Wilmad), with protein concentrations of 66 μM and 73 μM, respectively. enNTS_1_ΔM8A, ΔM7, ΔM6, ΔM5 spectra (red), and enNTS_1_ΔM8A (orange) were recorded at 800 MHz, in 5 mm Shigemi tubes, with protein concentrations of 20 μM, 20 μM, 21 μM, 24 μM, and 36 μM respectively.

**Supplemental Figure S7.**
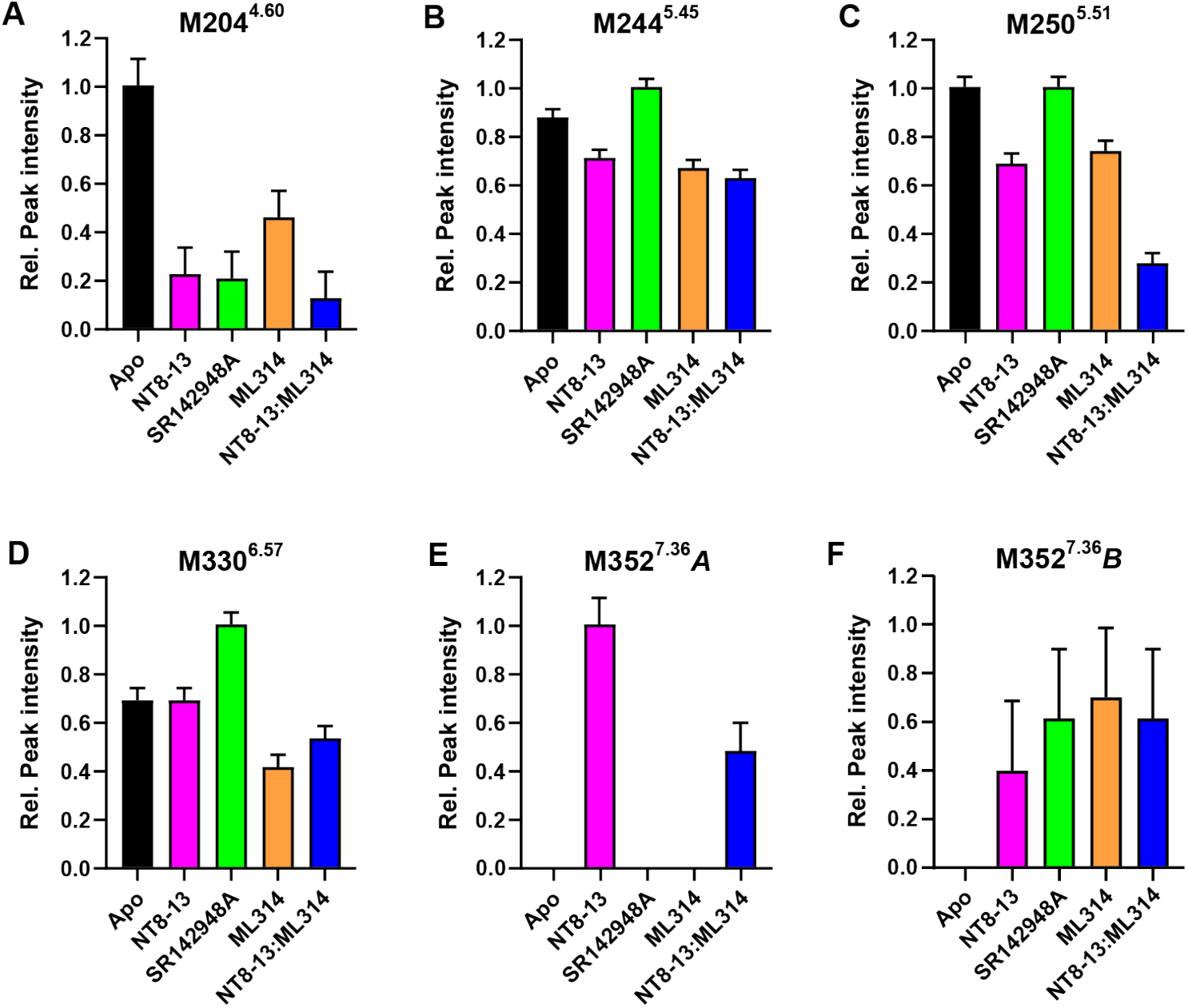
Ligands differentially affect enNTS_1_ΔM4 ^13^C^ε^H_3_-methionine peak intensities. Peak intensities were determined by manual integration using Sparky. For each residue, intensities were normalized to the largest valued condition. Bar color correspond to the color scheme used for HMQC spectra. M204^4.60^ intensities are the sum of all split resonances. Intensities for NT8-13 are the mean intensities from two independent samples. Error bars represent the standard deviation (SD) of peak intensities averaged for all 6 resonances in the NT8-13 duplicate experiments.

**Supplemental Figure S8.**
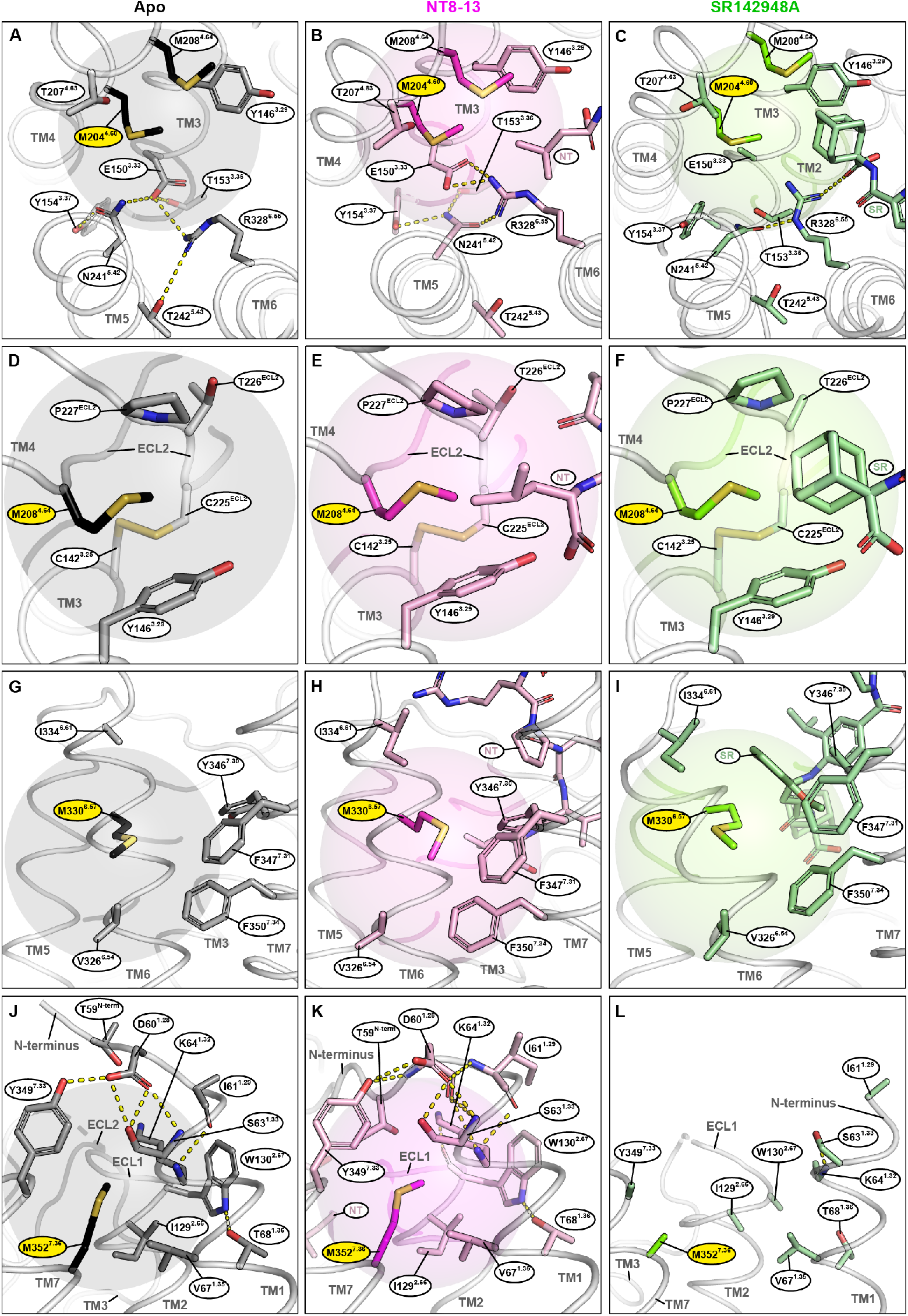
Chemical environments of NTS_1_ M204^4.60^, M208^4.64^, M330^6.57^, and M352^7.36^ observed in crystal structures. Transparent spheres indicate a 6 Å radius centred at the methionine C^ε^ atom, beyond which shielding effects of neighbouring amino acids are considered to be negligible ^3^. Yellow, dotted lines indicate hydrogen-bonds where relevant. A,D,G,J) Apo-state crystal structure (PDB 6Z66). B,E,H,K) NT8-13-bound crystal structure (PDB 6YVR, chain a). C,F,I,L) SR142948A-bound crystal structure (PDB 6Z4Q).

**Supplemental Figure S9.**
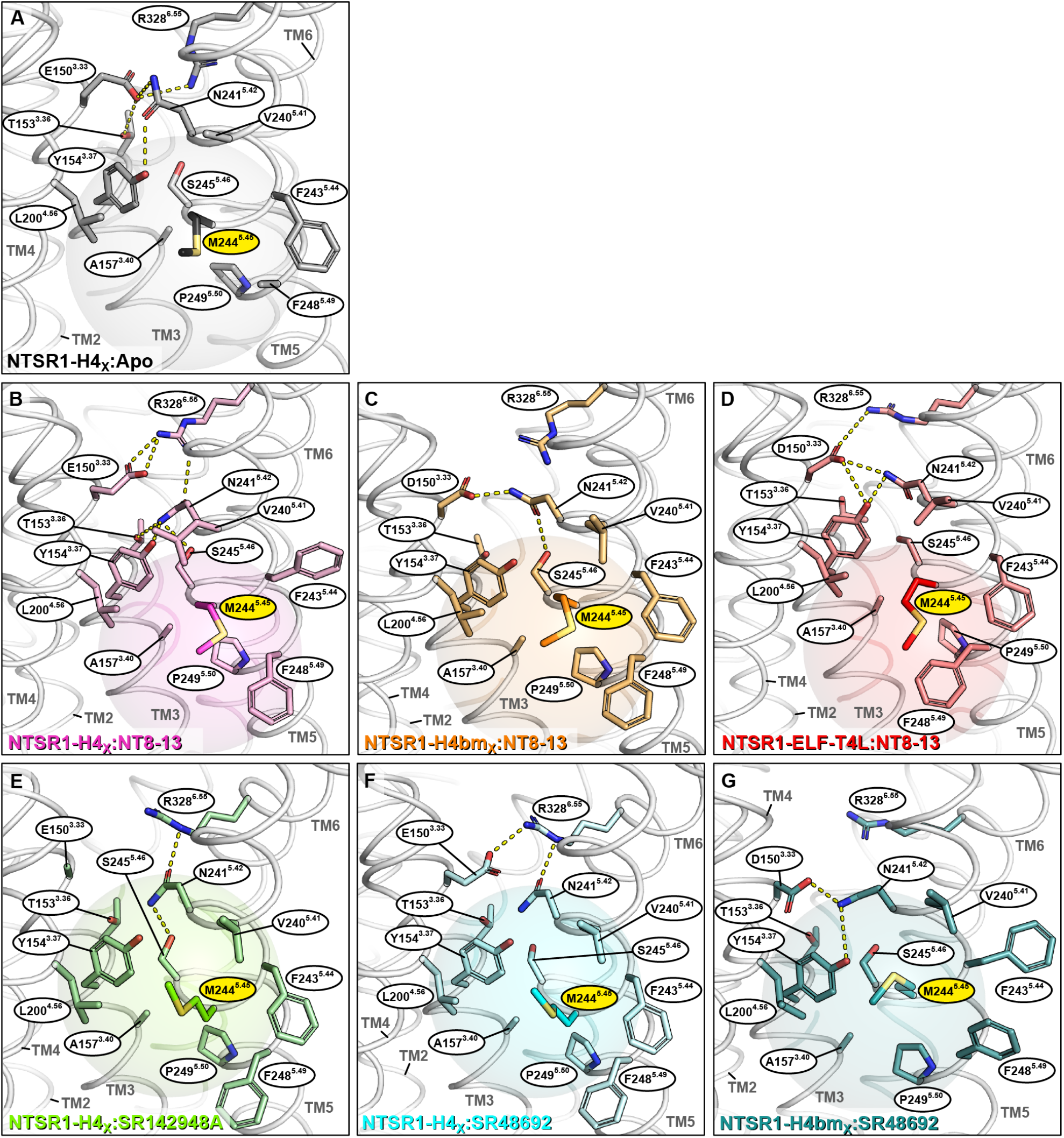
Chemical environments of NTS_1_ M244^5.45^ observed in crystal structures. Transparent spheres indicate a 6 Å radius centred at M244^5.45^C^ε^, beyond which shielding effects of neighbouring amino acids are considered to be negligible ^3^. Yellow, dotted lines indicate hydrogen-bonds where relevant. A) Apo-state crystal structure (PDB 6Z66); B) NT8-13-bound crystal structure (PDB 6YVR chain a); C) Back-mutated (BM) NT8-13-bound crystal structure (PDB 6Z4V); D) Active-like (ELF-T4L) NT8-13-bound crystal structure (PDB 4XEE); E) SR142948A-bound crystal structure (PDB 6Z4Q); F) SR48692-bound crystal structure (PDB 6ZIN); G) Back-mutated (BM) SR48692 crystal structure (PDB 6Z4S).

**Supplemental Figure S10.**
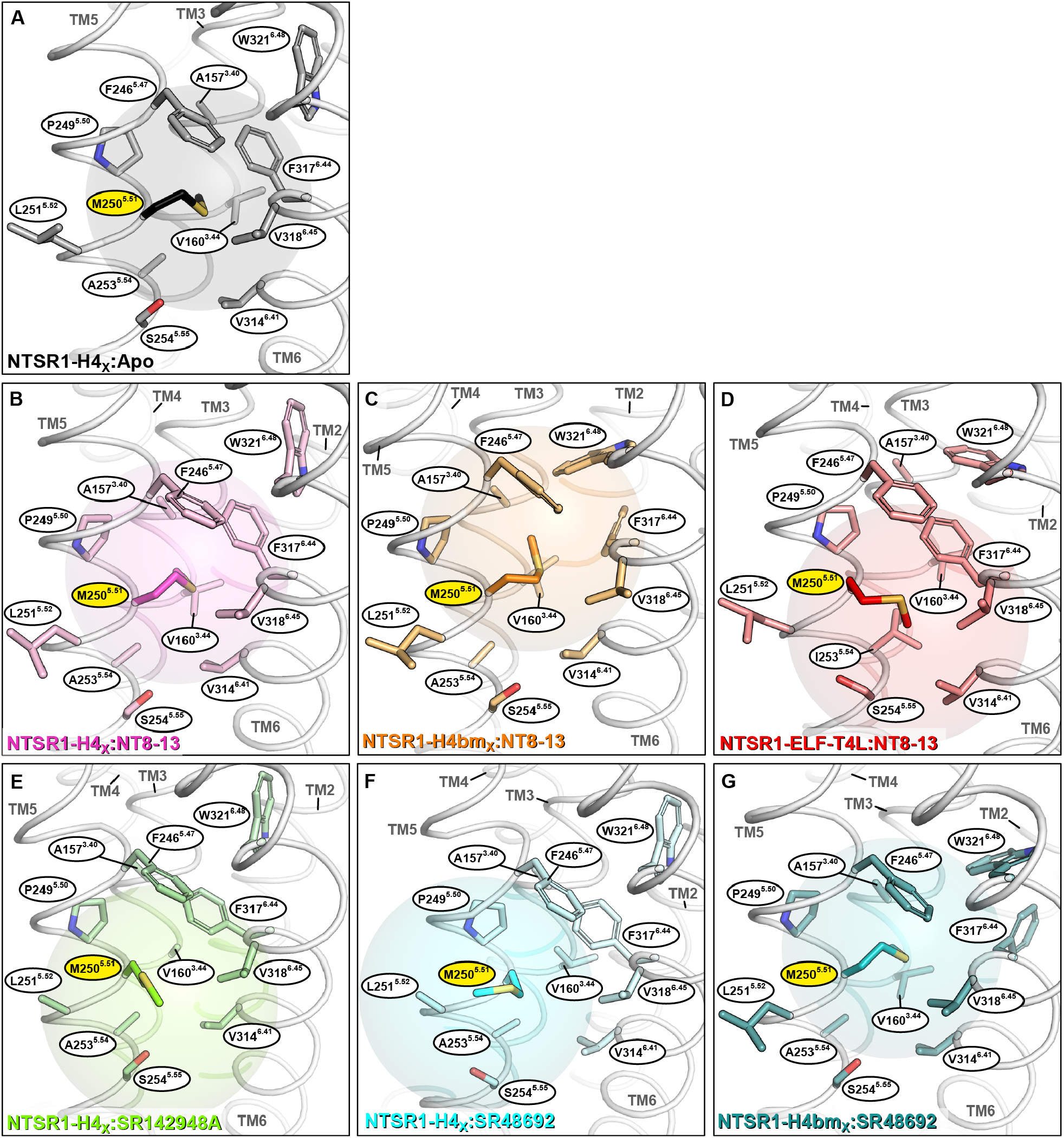
Chemical environments of NTS_1_ M250^5.51^ observed in crystal structures. Transparent spheres indicate a 6Å radius centred at M250^5.51^C^ε^, beyond which shielding effects of neighbouring amino acids are considered to be negligible ^3^. A) Apo-state crystal structure (PDB 6Z66); B) NT8-13-bound crystal structure (PDB 6YVR chain a); C) Back-mutated (BM) NT8-13-bound crystal structure (PDB 6Z4V); D) Active-like (ELF-T4L) NT8-13-bound crystal structure (PDB 4XEE); E) SR142948A-bound crystal structure (PDB 6Z4Q); F) SR48692-bound crystal structure (PDB 6ZIN); G) Back-mutated (BM) SR48692 crystal structure (PDB 6Z4S).

**Supplemental Figure S11.**
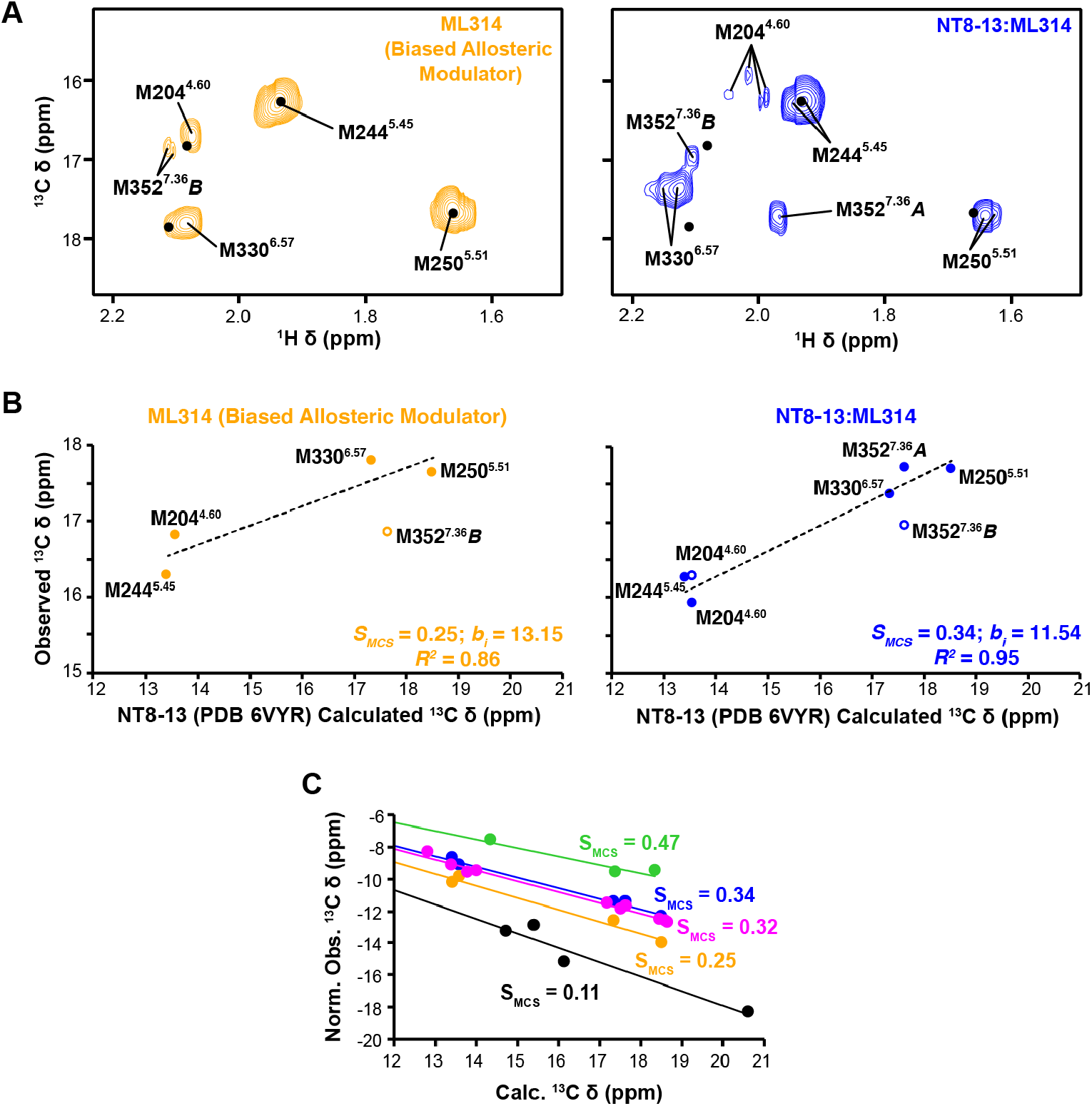
^1^H-^13^C HMQC spectra and global order parameter (S_MCS_) correlation plots for ML314:enNTS_1_ΔM4 and ML314:NT8-13:enNTS_1_ΔM4 complexes. A) ^1^H-^13^C HMQC spectra of [^13^C^ε^H_3_-methionine]-enNTS_1_ΔM4 in the presence of ML314 and NT8-13:ML314. Both spectra were recorded at 600 MHz with protein concentrations of 66 μM. B) Linear correlation between DFT-calculated and experimental ^13^C chemical shift values. The NT8-13:NTSR1-H4_X_ crystal structure (PDB 6VYR) was used as a “rigid-structure” reference. All observable resonances are included in the scatterplots. In spectra where a single methionine is assigned to multiple observable resonances, only the most populated (i.e. highest intensity) peak is fitted; the lower intensity peaks are typically outliers (open circle). C) Linear correlation plots of the normalized observed (minus DFT-calculated) and DFT-calculated chemical shift (Chashmniam et al., 2021). The slope of the normalized plots is S_MCS_-1 where S_MCS_ represents the global order parameters, with zero equal to completely averaged and one is completely rigid.

**Supplemental Figure S122.**
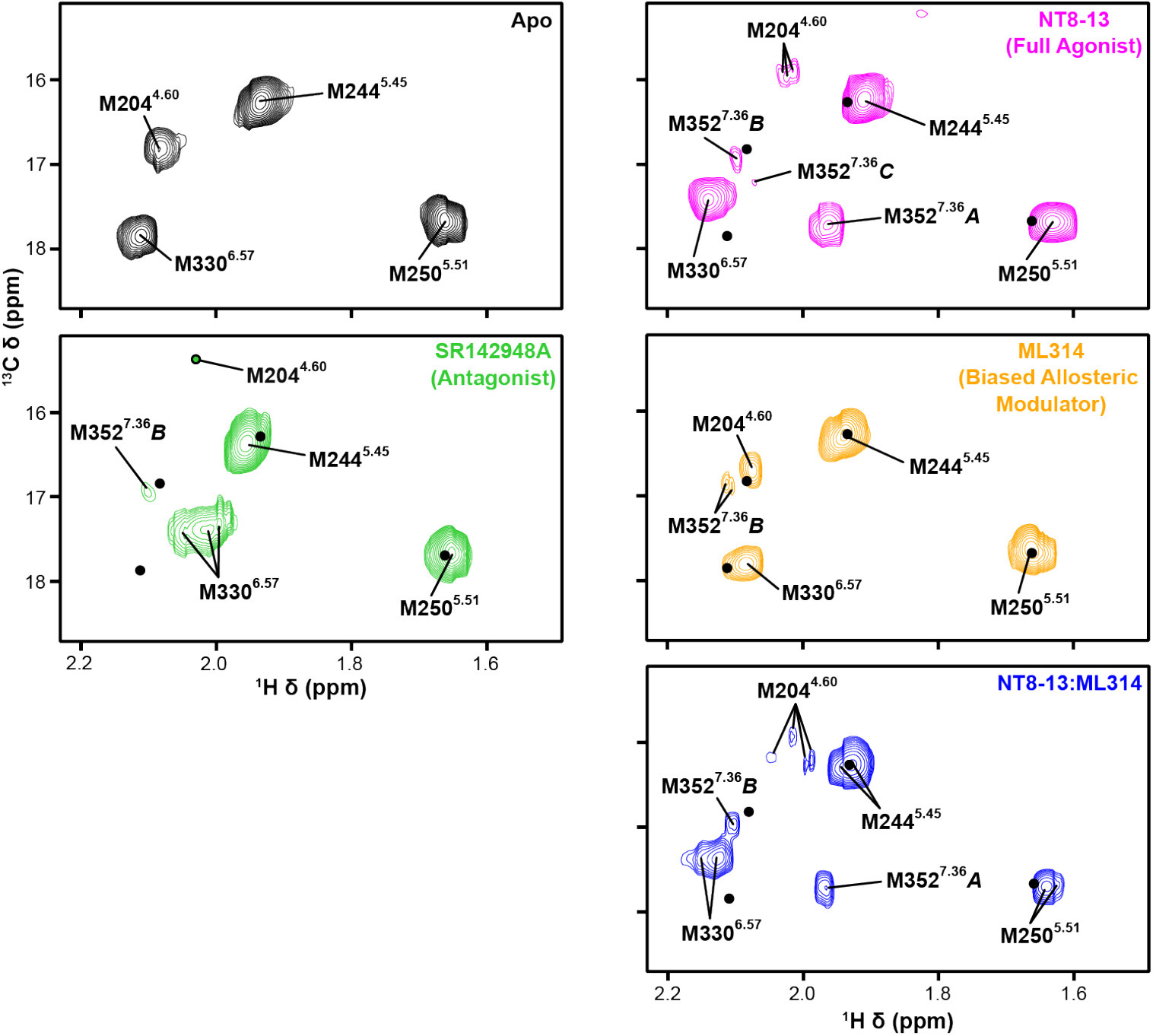
^1^H-^13^C HMQC spectra of enNTS_1_ΔM4 ^13^C^ε^H_3_-methionine for all ligand conditions. Colour coding of individual spectra match those used throughout the manuscript. All spectra were recorded at 600 MHz, in 3 mm thin wall precision NMR tubes (Wilmad), at enNTS_1_ΔM4 concentrations of 66 μM.

## LITERATURE CITED

1. Hauser, A. S.; Attwood, M. M.; Rask-Andersen, M.; Schioth, H. B.; Gloriam, D. E., Trends in GPCR drug discovery: new agents, targets and indications. Nat Rev Drug Discov 2017, 16 (12), 829–842.

2. Hauser, A. S.; Chavali, S.; Masuho, I.; Jahn, L. J.; Martemyanov, K. A.; Gloriam, D. E.; Babu, M. M., Pharmacogenomics of GPCR Drug Targets. Cell 2018, 172 (1-2), 41–54 e19.

3. Administration, U. F. a. D., Drug Trials Snapshots: OLINVYK. https://www.fda.gov/drugs/drug-approvals-and-databases/drug-trials-snapshots-olinvyk, 2020.

4. Administration, U. F. a. D., FDA Approves New Opioid for Intravenous Use in Hospitals, Other Controlled Clinical Settings. https://www.fda.gov/news-events/press-announcements/fda-approves-new-opioid-intravenous-use-hospitals-other-controlled-clinical-settings, 2020.

5. Kern, D.; Zuiderweg, E. R., The role of dynamics in allosteric regulation. Curr Opin Struct Biol 2003, 13 (6), 748–57.

6. Motlagh, H. N.; Wrabl, J. O.; Li, J.; Hilser, V. J., The ensemble nature of allostery. Nature 2014, 508 (7496), 331–9.

7. Thal, D. M.; Glukhova, A.; Sexton, P. M.; Christopoulos, A., Structural insights into G-protein-coupled receptor allostery. Nature 2018, 559 (7712), 45–53.

8. Lane, J. R.; May, L. T.; Parton, R. G.; Sexton, P. M.; Christopoulos, A., A kinetic view of GPCR allostery and biased agonism. Nat Chem Biol 2017, 13 (9), 929–937.

9. Rasmussen, S. G.; Choi, H. J.; Fung, J. J.; Pardon, E.; Casarosa, P.; Chae, P. S.; Devree, B. T.; Rosenbaum, D. M.; Thian, F. S.; Kobilka, T. S.; Schnapp, A.; Konetzki, I.; Sunahara, R. K.; Gellman, S. H.; Pautsch, A.; Steyaert, J.; Weis, W. I.; Kobilka, B. K., Structure of a nanobody-stabilized active state of the beta(2) adrenoceptor. Nature 2011, 469 (7329), 175–80.

10. Monod, J.; Wyman, J.; Changeux, J. P., On the Nature of Allosteric Transitions: A Plausible Model. Journal of molecular biology 1965, 12, 88–118.

11. Shimada, I.; Ueda, T.; Kofuku, Y.; Eddy, M. T.; Wuthrich, K., GPCR drug discovery: integrating solution NMR data with crystal and cryo-EM structures. Nat Rev Drug Discov 2018.

12. Cooper, A.; Dryden, D. T., Allostery without conformational change. A plausible model. Eur Biophys J 1984, 11 (2), 103–9.

13. Akke, M.; Brueschweiler, R.; Palmer, A. G., NMR order parameters and free energy: an analytical approach and its application to cooperative calcium(2+) binding by calbindin D9k. Journal of the American Chemical Society 1993, 115 (21), 9832–9833.

14. Caro, J. A.; Harpole, K. W.; Kasinath, V.; Lim, J.; Granja, J.; Valentine, K. G.; Sharp, K. A.; Wand, A. J., Entropy in molecular recognition by proteins. Proceedings of the National Academy of Sciences of the United States of America 2017, 114 (25), 6563–6568.

15. Kasinath, V.; Sharp, K. A.; Wand, A. J., Microscopic Insights into the NMR Relaxation-Based Protein Conformational Entropy Meter. Journal of the American Chemical Society 2013, 135 (40), 15092–15100.

16. Marlow, M. S.; Dogan, J.; Frederick, K. K.; Valentine, K. G.; Wand, A. J., The role of conformational entropy in molecular recognition by calmodulin. Nature Chemical Biology 2010, 6 (5), 352–8.

17. Hansen, D. F.; Neudecker, P.; Kay, L. E., Determination of Isoleucine Side-Chain Conformations in Ground and Excited States of Proteins from Chemical Shifts. Journal of the American Chemical Society 2010, 132 (22), 7589–7591.

18. London, R. E.; Wingad, B. D.; Mueller, G. A., Dependence of amino acid side chain 13C shifts on dihedral angle: application to conformational analysis. Journal of the American Chemical Society 2008, 130 (33), 11097–105.

19. Mulder, F. A. A., Leucine Side-Chain Conformation and Dynamics in Proteins from 13C NMR Chemical Shifts. ChemBioChem 2009, 10 (9), 1477–1479.

20. Chashmniam, S.; Teixeria, J. M. C.; Paniagua, J. C.; Pons, M., A methionine chemical shift based order parameter characterizing global protein dynamics. ChemBioChem 2021, 22 (6), 1001–1004.

21. Deluigi, M.; Klipp, A.; Klenk, C.; Merklinger, L.; Eberle, S. A.; Morstein, L.; Heine, P.; Mittl, P. R. E.; Ernst, P.; Kamenecka, T. M.; He, Y.; Vacca, S.; Egloff, P.; Honegger, A.; Plückthun, A., Complexes of the neurotensin receptor 1 with small-molecule ligands reveal structural determinants of full, partial, and inverse agonism. Science Advances 2021, 7 (5), eabe5504.

22. Bumbak, F.; Keen, A. C.; Gunn, N. J.; Gooley, P. R.; Bathgate, R. A. D.; Scott, D. J., Optimization and 13CH3 methionine labeling of a signaling competent neurotensin receptor 1 variant for NMR studies. Biochimica et Biophysica Acta (BBA) - Biomembranes 2018, 1860 (6), 1372–1383.

23. Ballesteros, J. A.; Weinstein, H., Integrated Methods for the Construction of Three-Dimensional Models and Computational Probing of Structure-Function Relations in G Protein-Coupled Receptors. Methods in Neurosciences 1995, 25.

24. Inoue, A.; Ishiguro, J.; Kitamura, H.; Arima, N.; Okutani, M.; Shuto, A.; Higashiyama, S.; Ohwada, T.; Arai, H.; Makide, K.; Aoki, J., TGFα shedding assay: an accurate and versatile method for detecting GPCR activation. Nature Methods 2012, 9, 1021.

25. Carraway, R.; Leeman, S. E., The amino acid sequence of a hypothalamic peptide, neurotensin. The Journal of biological chemistry 1975, 250 (5), 1907–11.

26. Barak, L. S.; Bai, Y.; Peterson, S.; Evron, T.; Urs, N.; Peddibhotla, S.; Hedrick, M. P.; Hershberger, P.; Maloney, P. R.; Chung, T. D. Y.; Rodriguiz, R. M.; Wetsel, W. C.; Thomas, J. B.; Hanson, G. R.; Pinkerton, A. B.; Caron, M. G., ML314: A Biased Neurotensin Receptor Ligand for Methamphetamine Abuse. ACS Chem Biol 2016.

27. Peddibhotla, S.; Hedrick, M. P.; Hershberger, P.; Maloney, P. R.; Li, Y. J.; Milewski, M.; Gosalia, P.; Gray, W.; Mehta, A.; Sugarman, E.; Hood, B.; Suyama, E.; Nguyen, K.; Heynen-Genel, S.; Vasile, S.; Salaniwal, S.; Stonich, D.; Su, Y.; Mangravita-Novo, A.; Vicchiarelli, M.; Roth, G. P.; Smith, L. H.; Chung, T. D. Y.; Hanson, G. R.; Thomas, J. B.; Caron, M. G.; Barak, L. S.; Pinkerton, A. B., Discovery of ML314, a Brain Penetrant Nonpeptidic beta-Arrestin Biased Agonist of the Neurotensin NTR1 Receptor. Acs Med Chem Lett 2013, 4 (9), 846–851.

28. Dixon, A. S.; Schwinn, M. K.; Hall, M. P.; Zimmerman, K.; Otto, P.; Lubben, T. H.; Butler, B. L.; Binkowski, B. F.; Machleidt, T.; Kirkland, T. A.; Wood, M. G.; Eggers, C. T.; Encell, L. P.; Wood, K. V., NanoLuc Complementation Reporter Optimized for Accurate Measurement of Protein Interactions in Cells. ACS Chem Biol 2016, 11 (2), 400–8.

29. Butterfoss, G. L.; DeRose, E. F.; Gabel, S. A.; Perera, L.; Krahn, J. M.; Mueller, G. A.; Zheng, X.; London, R. E., Conformational dependence of 13C shielding and coupling constants for methionine methyl groups. Journal of biomolecular NMR 2010, 48 (1), 31–47.

30. Mittermaier, A.; Kay, L. E.; Forman-Kay, J. D., Analysis of deuterium relaxation-derived methyl axis order parameters and correlation with local structure. Journal of biomolecular NMR 1999, 13 (2), 181–5.

31. Cong, X.; Fiorucci, S.; Golebiowski, J., Activation dynamics of the neurotensin G protein-coupled receptor 1. Journal of Chemical Theory and Computation 2018, 14 (8), 4467–4473.

32. Dror, R. O.; Arlow, D. H.; Maragakis, P.; Mildorf, T. J.; Pan, A. C.; Xu, H.; Borhani, D. W.; Shaw, D. E., Activation mechanism of the beta2-adrenergic receptor. Proceedings of the National Academy of Sciences of the United States of America 2011, 108 (46), 18684–9.

33. Baranac-Stojanović, M.; Koch, A.; Kleinpeter, E., Is the Conventional Interpretation of the Anisotropic Effects of CC Double Bonds and Aromatic Rings in NMR Spectra in Terms of the π-Electron Shielding/Deshielding Contributions Correct? Chemistry – A European Journal 2012, 18 (1), 370–376.

34. Labbejullie, C.; Botto, J. M.; Mas, M. V.; Chabry, J.; Mazella, J.; Vincent, J. P.; Gully, D.; Maffrand, J. P.; Kitabgi, P., [H-3]Sr-48692, the First Nonpeptide Neurotensin Antagonist Radioligand Characterization of Binding-Properties and Evidence for Distinct Agonist and Antagonist Binding Domains on the Rat Neurotensin Receptor. Mol Pharmacol 1995, 47 (5), 1050–1056.

35. Hilger, D., The role of structural dynamics in GPCR-mediated signaling. The FEBS journal 2021, 288 (8), 2461–2489.

36. Manglik, A.; Kim Tae H.; Masureel, M.; Altenbach, C.; Yang, Z.; Hilger, D.; Lerch Michael T.; Kobilka Tong S.; Thian Foon S.; Hubbell Wayne L.; Prosser, R. S.; Kobilka Brian K., Structural Insights into the Dynamic Process of β2-Adrenergic Receptor Signaling. Cell 2015, (0).

37. Wu, F.-J.; Williams, L. M.; Abdul-Ridha, A.; Gunatilaka, A.; Vaid, T. M.; Kocan, M.; Whitehead, A. R.; Griffin, M. D. W.; Bathgate, R. A. D.; Scott, D. J.; Gooley, P. R., Probing the correlation between ligand efficacy and conformational diversity at the α1A-adrenoreceptor reveals allosteric coupling of its microswitches. Journal of Biological Chemistry 2020.

38. Solt, A. S.; Bostock, M. J.; Shrestha, B.; Kumar, P.; Warne, T.; Tate, C. G.; Nietlispach, D., Insight into partial agonism by observing multiple equilibria for ligand-bound and Gs-mimetic nanobody-bound β1-adrenergic receptor. Nature communications 2017, 8 (1), 1795.

39. Kofuku, Y.; Ueda, T.; Okude, J.; Shiraishi, Y.; Kondo, K.; Maeda, M.; Tsujishita, H.; Shimada, I., Efficacy of the beta(2)-adrenergic receptor is determined by conformational equilibrium in the transmembrane region. Nature communications 2012, 3, 1045.

40. Kleckner, I. R.; Foster, M. P., An introduction to NMR-based approaches for measuring protein dynamics. Bba-Proteins Proteom 2011, 1814 (8), 942–968.

41. Barroso, S.; Richard, F.; Nicolas-Etheve, D.; Kitabgi, P.; Labbe-Jullie, C., Constitutive activation of the neurotensin receptor 1 by mutation of Phe(358) in Helix seven. Br J Pharmacol 2002, 135 (4), 997–1002.

42. Kling, R. C.; Plomer, M.; Lang, C.; Banerjee, A.; Hubner, H.; Gmeiner, P., Development of Covalent Ligand-Receptor Pairs to Study the Binding Properties of Nonpeptidic Neurotensin Receptor 1 Antagonists. ACS Chem Biol 2016, 11 (4), 869–75.

43. Frei, J. N.; Broadhurst, R. W.; Bostock, M. J.; Solt, A.; Jones, A. J. Y.; Gabriel, F.; Tandale, A.; Shrestha, B.; Nietlispach, D., Conformational plasticity of ligand-bound and ternary GPCR complexes studied by (19)F NMR of the beta1-adrenergic receptor. Nat Commun 2020, 11 (1), 669.

44. Isogai, S.; Deupi, X.; Opitz, C.; Heydenreich, F. M.; Tsai, C. J.; Brueckner, F.; Schertler, G. F.; Veprintsev, D. B.; Grzesiek, S., Backbone NMR reveals allosteric signal transduction networks in the beta1-adrenergic receptor. Nature 2016, 530 (7589), 237–41.

45. Kofuku, Y.; Ueda, T.; Okude, J.; Shiraishi, Y.; Kondo, K.; Mizumura, T.; Suzuki, S.; Shimada, I., Functional dynamics of deuterated beta2 -adrenergic receptor in lipid bilayers revealed by NMR spectroscopy. Angew Chem Int Ed Engl 2014, 53 (49), 13376–9.

46. Gully, D.; Labeeuw, B.; Boigegrain, R.; Oury-Donat, F.; Bachy, A.; Poncelet, M.; Steinberg, R.; Suaud-Chagny, M. F.; Santucci, V.; Vita, N.; Pecceu, F.; Labbe-Jullie, C.; Kitabgi, P.; Soubrie, P.; Le Fur, G.; Maffrand, J. P., Biochemical and pharmacological activities of SR 142948A, a new potent neurotensin receptor antagonist. The Journal of pharmacology and experimental therapeutics 1997, 280 (2), 802–12.

47. Kruse, A. C.; Ring, A. M.; Manglik, A.; Hu, J.; Hu, K.; Eitel, K.; Hubner, H.; Pardon, E.; Valant, C.; Sexton, P. M.; Christopoulos, A.; Felder, C. C.; Gmeiner, P.; Steyaert, J.; Weis, W. I.; Garcia, K. C.; Wess, J.; Kobilka, B. K., Activation and allosteric modulation of a muscarinic acetylcholine receptor. Nature 2013, advance online publication.

48. Sharp, K. A.; O’Brien, E.; Kasinath, V.; Wand, A. J., On the relationship between NMR-derived amide order parameters and protein backbone entropy changes. Proteins 2015, 83 (5), 922–30.

49. Leavitt, S.; Freire, E., Direct measurement of protein binding energetics by isothermal titration calorimetry. Curr Opin Struct Biol 2001, 11 (5), 560–6.

50. O’Connor, C.; White, K. L.; Doncescu, N.; Didenko, T.; Roth, B. L.; Czaplicki, G.; Stevens, R. C.; Wuthrich, K.; Milon, A., NMR structure and dynamics of the agonist dynorphin peptide bound to the human kappa opioid receptor. Proceedings of the National Academy of Sciences of the United States of America 2015, 112 (38), 11852–7.

51. Okude, J.; Ueda, T.; Kofuku, Y.; Sato, M.; Nobuyama, N.; Kondo, K.; Shiraishi, Y.; Mizumura, T.; Onishi, K.; Natsume, M.; Maeda, M.; Tsujishita, H.; Kuranaga, T.; Inoue, M.; Shimada, I., Identification of a Conformational Equilibrium That Determines the Efficacy and Functional Selectivity of the μ-Opioid Receptor. Angewandte Chemie International Edition 2015, 54, 15771–15776.

52. Xu, J.; Hu, Y.; Kaindl, J.; Risel, P.; Hübner, H.; Maeda, S.; Niu, X.; Li, H.; Gmeiner, P.; Jin, C.; Kobilka, B. K., Conformational Complexity and Dynamics in a Muscarinic Receptor Revealed by NMR Spectroscopy. Molecular Cell 2019.

53. Lipari, G.; Szabo, A., Model-free approach to the interpretation of nuclear magnetic resonance relaxation in macromolecules. 1. Theory and range of validity. Journal of the American Chemical Society 1982, 104 (17), 4546–4559.

54. Lipari, G.; Szabo, A., Model-free approach to the interpretation of nuclear magnetic resonance relaxation in macromolecules. 2. Analysis of experimental results. Journal of the American Chemical Society 1982, 104 (17), 4559–4570.

55. Tzeng, S. R.; Kalodimos, C. G., Protein activity regulation by conformational entropy. Nature 2012, 488 (7410), 236–40.

56. Wand, A. J.; Sharp, K. A., Measuring Entropy in Molecular Recognition by Proteins. Annu Rev Biophys 2018, 47, 41–61.

57. Goto, N. K.; Gardner, K. H.; Mueller, G. A.; Willis, R. C.; Kay, L. E., A robust and cost-effective method for the production of Val, Leu, Ile (delta 1) methyl-protonated 15N-, 13C-, 2H-labeled proteins. Journal of biomolecular NMR 1999, 13 (4), 369–74.

58. Ishima, R.; Louis, J. M.; Torchia, D. A., Optimized labeling of 13CHD2 methyl isotopomers in perdeuterated proteins: potential advantages for 13C relaxation studies of methyl dynamics of larger proteins. Journal of biomolecular NMR 2001, 21 (2), 167–71.

59. Sun, H.; Kay, L. E.; Tugarinov, V., An optimized relaxation-based coherence transfer NMR experiment for the measurement of side-chain order in methyl-protonated, highly deuterated proteins. J Phys Chem B 2011, 115 (49), 14878–84.

60. Linser, R.; Gelev, V.; Hagn, F.; Arthanari, H.; Hyberts, S. G.; Wagner, G., Selective methyl labeling of eukaryotic membrane proteins using cell-free expression. Journal of the American Chemical Society 2014, 136 (32), 11308–10.

61. Opitz, C.; Isogai, S.; Grzesiek, S., An economic approach to efficient isotope labeling in insect cells using homemade 15N-, 13C- and 2H-labeled yeast extracts. Journal of biomolecular NMR 2015, 62 (3), 373–85.

62. Sitarska, A.; Skora, L.; Klopp, J.; Roest, S.; Fernandez, C.; Shrestha, B.; Gossert, A. D., Affordable uniform isotope labeling with (2)H, (13)C and (15)N in insect cells. Journal of biomolecular NMR 2015, 62 (2), 191–7.

63. Kerfah, R.; Plevin, M. J.; Sounier, R.; Gans, P.; Boisbouvier, J., Methyl-specific isotopic labeling: a molecular tool box for solution NMR studies of large proteins. Curr Opin Struct Biol 2015, 32, 113–22.

64. Luginbühl, P.; Wüthrich, K., Semi-classical nuclear spin relaxation theory revisited for use with biological macromolecules. Progress in Nuclear Magnetic Resonance Spectroscopy 2002, 40 (3), 199–247.

65. Schanda, P.; Brutscher, B., Very fast two-dimensional NMR spectroscopy for real-time investigation of dynamic events in proteins on the time scale of seconds. Journal of the American Chemical Society 2005, 127 (22), 8014–5.

66. Bumbak, F.; Bathgate, R. A. D.; Scott, D. J.; Gooley, P. R., Expression and Purification of a Functional E. coli 13CH3-Methionine-Labeled Thermostable Neurotensin Receptor 1 Variant for Solution NMR Studies. In G Protein-Coupled Receptor Signaling: Methods and Protocols, Tiberi, M., Ed. Springer New York: New York, NY, 2019; pp 31–55.

67. Dixon, A. D.; Inoue, A.; Robson, S. A.; Culhane, K. J.; Trinidad, J. C.; Sivaramakrishnan, S.; Bumbak, F.; Ziarek, J. J., Effect of Ligands and Transducers on the Neurotensin Receptor 1 Conformational Ensemble. Journal of the American Chemical Society 2022, 144 (23), 10241–10250.

68. Scott, D. J.; Gunn, N. J.; Yong, K. J.; Wimmer, V. C.; Veldhuis, N. A.; Challis, L. M.; Haidar, M.; Petrou, S.; Bathgate, R. A. D.; Griffin, M. D. W., A Novel Ultra-Stable, Monomeric Green Fluorescent Protein For Direct Volumetric Imaging of Whole Organs Using CLARITY. Scientific Reports 2018, 8 (1), 667.

69. Sarkar, C. A.; Dodevski, I.; Kenig, M.; Dudli, S.; Mohr, A.; Hermans, E.; Pluckthun, A., Directed evolution of a G protein-coupled receptor for expression, stability, and binding selectivity. Proceedings of the National Academy of Sciences of the United States of America 2008, 105 (39), 14808–13.

70. Zhao, H. M.; Zha, W. J., In vitro ‘sexual’ evolution through the PCR-based staggered extension process (StEP). Nat Protoc 2006, 1 (4), 1865–1871.

71. Kupce, E.; Freeman, R., Polychromatic Selective Pulses. Journal of Magnetic Resonance, Series A 1993, 102 (1), 122–126.

72. Kupce, E.; Boyd, J.; Campbell, I. D., Short Selective Pulses for Biochemical Applications. Journal of Magnetic Resonance, Series B 1995, 106 (3), 300–303.

73. Kazimierczuk, K.; Orekhov, V. Y., Accelerated NMR spectroscopy by using compressed sensing. Angew Chem Int Ed Engl 2011, 50 (24), 5556–9.

74. Delaglio, F.; Grzesiek, S.; Vuister, G. W.; Zhu, G.; Pfeifer, J.; Bax, A., NMRPipe: a multidimensional spectral processing system based on UNIX pipes. Journal of biomolecular NMR 1995, 6 (3), 277–93.

75. Shihoya, W.; Izume, T.; Inoue, A.; Yamashita, K.; Kadji, F. M. N.; Hirata, K.; Aoki, J.; Nishizawa, T.; Nureki, O., Crystal structures of human ET(B) receptor provide mechanistic insight into receptor activation and partial activation. Nat Commun 2018, 9 (1), 4711.

76. Krumm, B. E.; White, J. F.; Shah, P.; Grisshammer, R., Structural prerequisites for G-protein activation by the neurotensin receptor. Nature communications 2015, 6, 7895.

## References

1. Inoue, A.; Ishiguro, J.; Kitamura, H.; Arima, N.; Okutani, M.; Shuto, A.; Higashiyama, S.; Ohwada, T.; Arai, H.; Makide, K.; Aoki, J., TGFα shedding assay: an accurate and versatile method for detecting GPCR activation. Nature Methods 2012, 9, 1021.

2. Dixon, A. S.; Schwinn, M. K.; Hall, M. P.; Zimmerman, K.; Otto, P.; Lubben, T. H.; Butler, B. L.; Binkowski, B. F.; Machleidt, T.; Kirkland, T. A.; Wood, M. G.; Eggers, C. T.; Encell, L. P.; Wood, K. V., NanoLuc Complementation Reporter Optimized for Accurate Measurement of Protein Interactions in Cells. ACS Chemical Biology 2016, 11 (2), 400-408.

3. Chashmniam, S.; Teixeria, J. M. C.; Paniagua, J. C.; Pons, M., A methionine chemical shift based order parameter characterizing global protein dynamics. Chembiochem 2020

